# Homotopic contralesional excitation suppresses spontaneous circuit repair and global network reconnections following ischemic stroke

**DOI:** 10.1101/2021.05.02.442355

**Authors:** Annie R. Bice, Qingli Xiao, Justin Kong, Ping Yan, Zachary P. Rosenthal, Andrew W. Kraft, Karen Smith, Tadeusz Wieloch, Jin-Moo Lee, Joseph P. Culver, Adam Q. Bauer

## Abstract

Understanding circuit-level changes that affect the brain’s capacity for plasticity will inform the design of targeted interventions for treating stroke recovery. We combine optogenetic photostimulation with optical neuroimaging to examine how contralesional excitatory activity affects cortical remodeling after stroke in mice. Following photothrombosis of left primary somatosensory forepaw (S1FP) cortex, mice received chronic excitation of right S1FP, a maneuver mimicking the use of the unaffected limb during recovery. Contralesional excitation suppressed perilesional S1FP remapping and was associated with abnormal patterns of evoked activity in the unaffected limb. Contralesional stimulation prevented the restoration of resting-state functional connectivity (RSFC) within the S1FP network, RSFC in several networks functionally-distinct from somatomotor regions, and resulted in persistent limb-use asymmetry. In stimulated mice, perilesional tissue exhibited suppressed transcriptional changes in several genes important for recovery. These results suggest that contralesional excitation impedes local and global circuit reconnection through suppression of several neuroplasticity-related genes after stroke.

## Introduction

Stroke causes direct structural damage to local brain circuits and indirect functional damage to global networks resulting in behavioral deficits spanning multiple domains[1, 2]. In the months following stroke, functional magnetic resonance imaging (FMRI) studies have shown that local circuits lost to infarction remodel in periinfarct cortex[3]. This process, termed “remapping”, appears to be tightly focused to perilesional regions in patients exhibiting good recovery, and suggests that periinfarct cortex may take over the function of brain regions lost to stroke [4–6]. Similar studies examining brain-wide patterns of synchronized, resting-state, hemodynamic activity after stroke have shown that global patterns of resting-state functional connectivity (RSFC) are also altered [7, 8]. Shortly after ischemic stroke, disruption of interhemispheric homotopic RSFC predicts poor motor and attentional recovery [9, 10], and in rats, restoration of homotopic RS-FC correlates with improved behavioral performance [11]. Similarly, we have shown in mice that functional disruption correlates with infarct size, regional RSFC disruption, the degree of remapping, and behavior [12, 13]. While functional neuroimaging studies consistently demonstrate local and global changes in functional brain organization after stroke, it is unknown how these processes interrelate to support recovery of function.

Activity in brain regions functionally-connected to perilesional tissue can differentially affect recovery potential. In monkeys, the use of the affected limb is required for remapping [6], and conversely, overactivation of the contralateral hemisphere through the use of the “good limb” is associated with poorer clinical outcome [14, 15]. Additionally, there is substantial indirect evidence that homotopic, interhemispheric connections may directly impact recovery after stroke. Stroke patients with impaired hand movement experience abnormally high interhemispheric inhibition exerted from intact contralesional motor cortex to perilesional motor cortex, with the degree of inhibition increasing with deficit severity[16]. It is thought that an equilibrium between excitation/inhibition across hemispheres may be important for normal unilateral function[17]. If the interhemispheric balance is disrupted (e.g. by stroke) local ipsilesional inhibitory influences from distant connections may worsen functional impairment. Prior human studies have primarily used noninvasive brain stimulation techniques (e.g., repetitive transcranial magnetic stimulation (rTMS)) to investigate the role of contralateral activity on the brain[18]. However, treatment efficacy using these methods is extremely varied [19-22], and are limited by imprecision and indiscriminate activation or inhibition of all cell types near the stimulated site. Thus, it has been difficult to systematically examine the influence of specific connection pathways important for recovery in human stroke patients [16, 23, 24]. Optogenetics is a powerful approach for increasing the specificity of paradigms designed to modulate local and distant neural activity after experimental stroke [25–29]. The aim of this study was to determine if local circuits lost due to stroke in mice remap into perilesional cortex and reconnect with global functional circuits (i.e. resting state networks), and if this process could be manipulated (spatially and temporally) by contralesional, homotopic excitation. Our results suggest that limited, but focal excitation of contralesional cortex impedes perilesional remapping and restoration of RSFC within the somatomotor network. Further, this maneuver profoundly altered the brain’s global functional connectome post stroke, and was associated with suppression of several molecular pathways important for recovery after injury.

## Results

### Study Design

In this study, we used optogenetic targeting in conjunction with optical intrinsic signal imaging (OISI) in mice to examine the effects of contralesional excitation on local and global cortical remodeling after focal ischemia. Our experimental design is illustrated in **Figure 1**. Note that all cortical regions and manipulations are in reference to hemisphere (instead of limb to which a region is mapped). We performed photothrombosis targeted to left primary somatosensory forepaw (S1FP) cortex. A random subset of mice was selected to receive chronic, intermittent contralesional excitation of right S1FP (+Stim), a maneuver mimicking the use of the unaffected limb and thought to negatively impact functional recovery after stroke. Contralesional excitation (3 min/day total) was delivered beginning day 1 after photothrombosis and continued for 5 consecutive days/week over 4 weeks. The remaining mice recovered spontaneously (−Stim). Assessments of functional recovery (via cylinder rearing) and OISI of stimulus evoked responses and RSFC occurred before, and 1 and 4 weeks after stroke under Ketamine/Xylazine anesthesia. Infarct volume and mRNA expression were characterized in a subset of mice from both stroke groups on days 5 and 14 post stroke, respectively. A sham group controlled for the effects of chronic photostimulation of Right S1FP cortex in the absence of stroke and were imaged at the same time points as the stroke groups.

**Figure 1.**
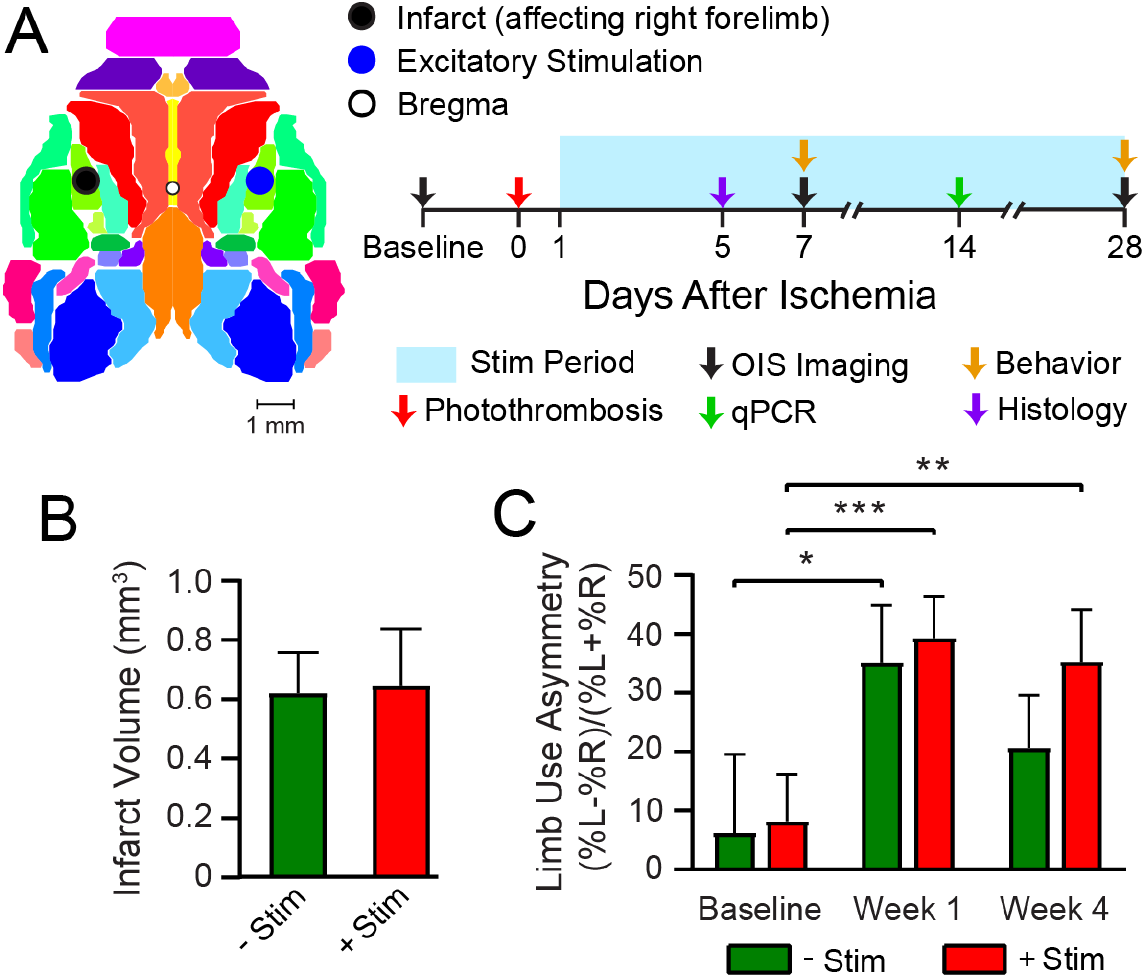
Graphical illustration of experiment and timeline. A) Photothrombosis was delivered to primary somatosensory forepaw cortex (S1FP) in the left hemisphere. Beginning on day 1 after focal ischemia, a subset of mice were subjected to chronic, intermittent optogenetic photostimulation of homotopic S1FP in the right hemisphere. Treatment was given for 5 consecutive days/week for 4 weeks. Optical intrinsic signal imaging of stimulus evoked and resting state activity occurred before, and 1 and 4 weeks after stroke. Infarct Volume and mRNA expression were characterized in a subset of mice from both groups at 5 and 14 days after photothrombosis. B) Infarct volume characterized 5 days post photothrombosis for −Stim (n=5) and+Stim (n=4) groups were statistically equivalent. C) Limb use as measured by cylinder rearing test in −Stim (n=15) and +Stim (n=20) groups. Symmetrical limb use was observed at baseline time points in all mice. Photothrombosis resulted in use asymmetry due to decreased use of right forelimb within the first week of both groups. – Stim mice demonstrated significant improvement by week 4 while+Stim mice exhibited sustained use asymmetry at 4 weeks. Repeated measures 2-way ANOVA revealed a main effect of Time (p=0.0004). Post hoc tests were performed as t-tests assuming unequal group variance and corrected for multiple comparisons using false discovery rate correction. *=p<0.05; **=p<0.01; ***=p<0.001 compared to baseline time-point. All data reported as mean ± S.E.

### Contralesional excitation inhibits behavioral recovery and cortical remapping and after stroke

We implemented a photostimulation protocol previously reported to positively affect recovery when chronically applied to ipsilesional tissue after stroke[25]. To ensure that our stimulation paradigm did not exacerbate the degree of initial injury, average infarct volume (in mm^3^) was quantified 5 days after photothrombosis. Both groups had statistically equivalent lesion sizes (p=0.92, **Fig. 1B**). Using the cylinder rearing test, symmetrical limb use was observed at baseline in all mice (**Fig. 1C**), with no group differences before stroke (p=0.9). Photothrombosis resulted in decreased use of right forelimb within the first week of both groups (−Stim mice: 35%, p=0.018 and +Stim mice: 39%, p=0.001). Similar performance deficits were observed in both groups at week 1 (p=0.73). At week 4, performance in –Stim mice was comparable to baseline (p=0.29), while +Stim mice continued to exhibit sustained use asymmetry compared to baseline (p=0.014). Limb use asymmetry did not significantly differ between groups at week 4 (p=0.25).

Recovery of sensorimotor function is associated with the formation of new cortical representations of the affected modality (remapping)[12, 30]. Because use of the unaffected limb is thought to be detrimental to recovery, we hypothesized that contralesional excitation would negatively impact remapping (explored in **Figs. 2, 3**). At baseline, electrical stimulation of the right forepaw (affected limb) resulted in consistent activation of left S1FP cortex in 100% of mice in both groups (**Figs. 2A, 3A**). Photothrombosis targeted to left S1FP resulted in >70% loss of evoked response magnitude in the first week following injury in both groups (p<0.001), and at least an 80% reduction in response area in both groups (p<0.0001, **Fig. 2B**). Responses from the affected limb reappeared in 80% of the mice in the –Stim group (**Fig. 3A**) by 4 weeks, and exhibited magnitudes statistically indistinguishable (p=0.22) from baseline responses (**Fig. 2B**). These new cortical representations in –Stim mice were focused to regions lateral and posterior to the infarct (near barrel cortex) (**Fig. 2A, Fig. 3A**), significantly larger than those at week 1 (p=0.045), but remained smaller than baseline representations (p=0.0002, **Fig. 2B**). In +Stim mice, only 67% of mice had detectible responses at week 4 (**Fig. 3A**). Week 4 responses in +Stim mice were 60% smaller in magnitude (p=0.0004) and 87% smaller in area (p<0.0001) than those at baseline and were indistinguishable from response area (p=0.92) and magnitude (p=0.63) observed at week 1 (**Fig. 2B**).

**Figure 2.**
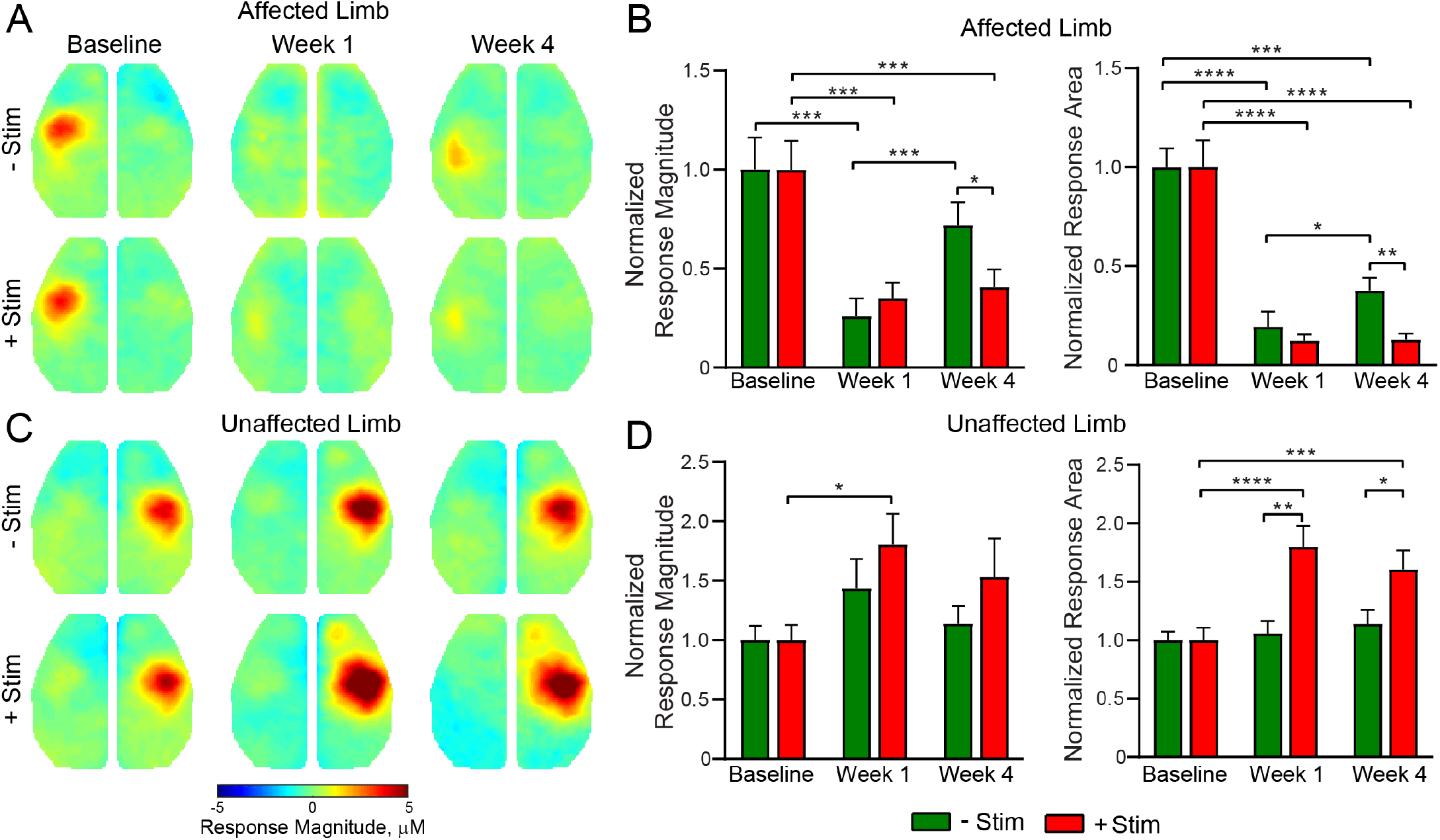
Contralesional activity inhibits cortical remapping after stroke. A) In the affected (right) limb, stimulus evoked responses following electrical forepaw stimulation were significantly reduced 1 week after stroke in both groups. Mice recovering spontaneously (−Stim, n=10, top row) exhibited robust activations at 4 weeks, while responses in stimulated mice (+Stim, n=15) were comparable to week 1 responses. B) Quantification of evoked response magnitude and area in the ipsilesional hemisphere. Repeated measures 2-way ANOVA revealed a significant Group x Time interaction (p=0.033) for evoked response magnitude and a trending Group x Time interaction (p=0.057) in response area. Main effects of Group (p=0.005) and Time (0.045) were observed for response area while magnitude also exhibited a main effect of Time (p=0.008). C) In the unaffected (left) limb, −Stim mice exhibited a trend towards increased activity at Week 1 that subsided by Week 4. In +Stim mice, evoked responses were significantly elevated at Week 1 and 4 compared to baseline. D) Quantification of evoked response magnitude and area in the contralesional hemisphere. Repeated measures 2-way ANOVA revealed a significant Group x Time interaction (p=0.0048), as well as a main effect of Time (p=0.0009) and Group (p=0.02) for evoked response area. A main effect of Time (p=0.0051) was observed for response magnitude. Post hoc tests were performed as t-tests assuming unequal group variance and corrected for multiple comparisons using false discovery rate correction. All analysis performed using total hemoglobin as contrast. *=p<0.05; **=p<0.01; ***=p<0.001; ****=p<0.0001.

**Figure 3.**
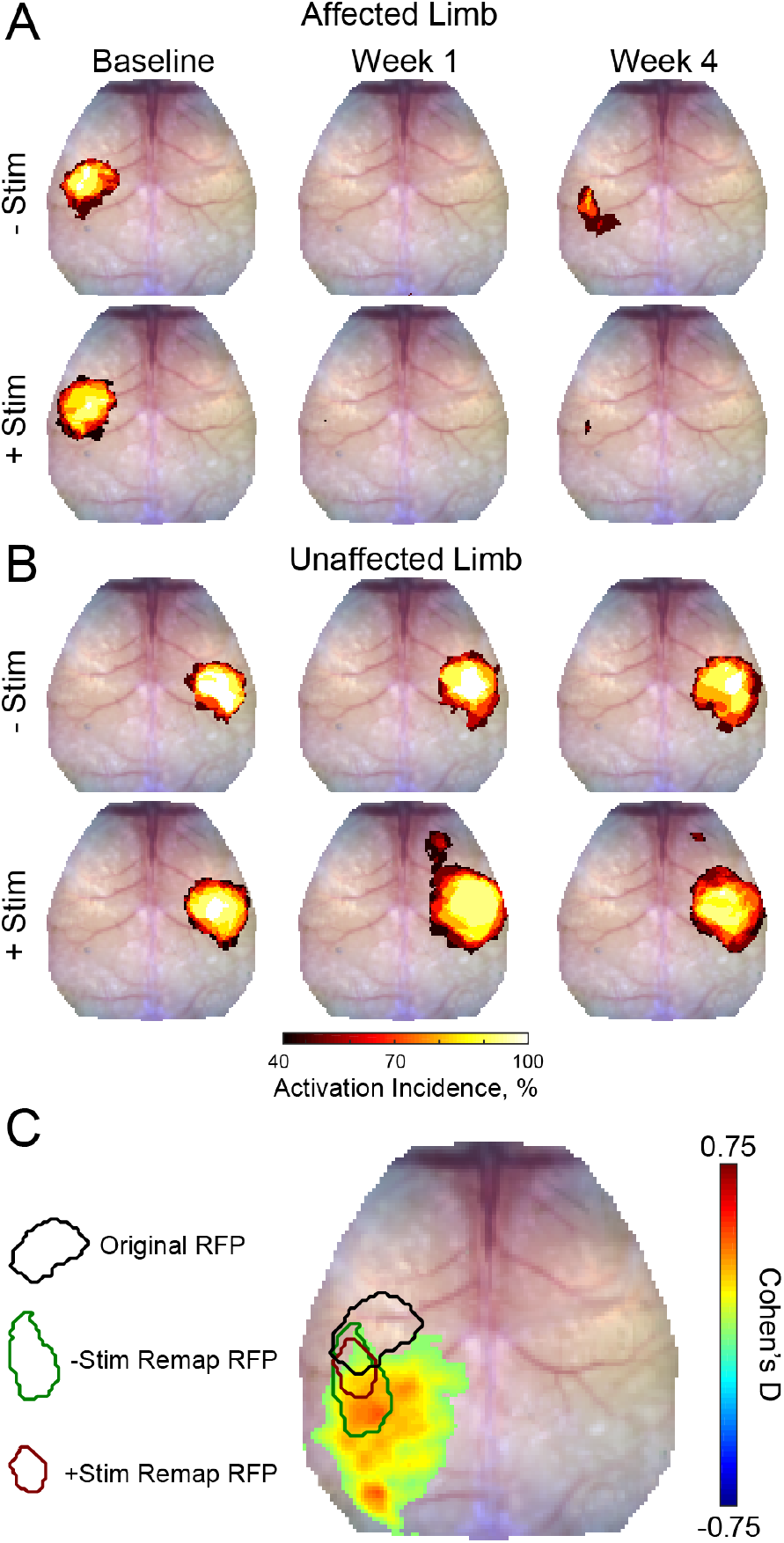
Activation incidence and effect size of forepaw remapping. Incidence of evoked responses for each group in the A) Affected and B) Unaffected limb (−Stim (n=10) and +Stim (n=15)). 100% of mice produced robust responses at baseline in both limbs. 1 week after stroke both groups exhibited significantly reduced activations in the affected limb, with at most 50% of mice having detectible responses in limited regions of cortex. 4 weeks after stroke, 80% of − Stim mice exhibited reproducible responses in large regions of somatosensory cortex, posterior to the original forepaw region. Conversely, only 67% of +Stim mice had detectible responses that were smaller in total area. B) In the unaffected limb, responses in the –Stim mice were largely similar across time points. However, responses in +Stim mice were generally larger (in line with Figure 2) and exhibited increased activity in forepaw motor cortex compared to the baseline time point. C) Cohen’s D was calculated as –Stim minus +Stim so that an effect would manifest as a positive value. Main topographical difference between groups included the region of forepaw remapping in the –Stim group. Contours of original and remapped forepaw response shown for reference.

Left forepaw (unaffected limb) stimulation resulted in consistent activation of right S1FP cortex in 100% of the mice in both groups prior to photothrombosis (**Fig. 2C, Fig 3B)**. In +Stim mice, evoked responses from the unaffected limb at week 1 were 81% larger than baseline responses (p=0.003, **Fig. 2D**). Activation magnitude in −Stim mice trended (p=0.1) towards larger responses at week 1, but returned to baseline levels by week 4 (p=0.2). In +Stim mice, activation area was 80% larger at week 1 (p<0.0001) and 60% larger at week 4 (p=0.001) than those at baseline. Further, stimulation of the unaffected limb in+Stim mice activated forepaw representations in contralesional motor cortex in 80% of mice that were not observed at baseline (**Fig. 3B**). Post stroke activation area in −Stim mice did not significantly differ from baseline at either time point (Week 1: p=0.66; Week 4: 0.21, **Fig. 2D**), nor elicit responses in contralesional motor cortex (**Fig. 3B**).

To further examine the effect of contralesional stimulation on group-wise cortical remapping, T-maps were calculated for each mouse for all imaging sessions of evoked activity at 4 weeks. From these T-maps, a map of Cohen’s D was calculated as –Stim minus +Stim so that an effect would manifest as a positive value (**Fig. 3C**). Computed differences between groups included the region of forepaw remapping in the ipsilesional hemisphere dominated by −Stim mice. Peak Cohen’s D in the region of remapping was 0.44, an intermediate effect, indicating that −Stim mice exhibit a larger, more active portion of remapped cortex compared to +Stim mice.

### Focal excitatory stimulation in control mice results in increased evoked responses of right and left S1FP

To determine if the effects of contralesional stimulation were conditional on focal ischemia, stimulus evoked responses were characterized in mice experiencing chronic excitatory stimulation of right S1FP in the absence of photothrombosis (**Fig. S1**). Robust evoked activity was observed in all control mice and time points (**Fig. S1A**). Interestingly, by 4 weeks, this group also exhibited increased evoked activity in forepaw representations of motor cortex, similar to contralesional responses observed in +Stim mice after stroke. In control mice, stimulus evoked responses in the right hemisphere exhibited larger response area at 1 week (p=0.002) and 4 weeks (p=0.0003) compared to baseline, and between 1 week and 4 weeks (p=0.004) (**Fig. S1B** top). Response magnitude in the right hemisphere was larger at week 4 compared to week 1 (p=0.01) (**Fig. S1B** top). In the left hemisphere, response area was significantly increased at 4 weeks compared to baseline (p=0.004) and at 4 weeks compared to 1 week (p=0.02) (**Fig. S1B** bottom). Response magnitude did not significantly change over time, but trended towards a larger response between week 1 and 4 (uncorrected p=0.037).

### Contralesional stimulation inhibits recovery of somatomotor forepaw circuitry

Both remapping[4, 12] and the formation of new homotopic functional connections[9, 11] are associated with better recovery following stroke. Having observed differential remapping in +Stim and –Stim groups, we examined how contralesional excitation affected reintegration of local S1FP circuitry into the somatomotor resting-state network. ROIs for RSFC analysis were defined by stimulus evoked responses at baseline and week 4 (see Methods): Original S1FP representations in both hemispheres, new (remapped) S1FP representations in the lesioned hemisphere, and forepaw representations in contralesional motor cortex, M1FP, (**Fig. 4A**). Group-averaged S1FP RSFC maps at baseline reveal positive ipsilateral RSFC with sensory and motor regions, as well as bilateral RSFC with original S1FP in both groups (**Fig. 4B**). 1 week after photothrombosis, bilateral RSFC between original ipsilesional and contralesional S1FP in both groups is significantly reduced (−Stim: p=0.0002; +Stim: p=0.008) (**Fig. 4B,C**). Further, at week 1, RSFC between contralesional S1FP and the region of future ipsilesional remapping has not been established in either group (- Stim: p=0.0014; +Stim: 0.0036)(**Fig. 4B, D, Fig. S2A**). However, in −Stim mice 4 weeks after stroke new “homotopic” functional connections between contralesional S1FP and remapped ipsilesional S1FP were observed having RSFC strength equivalent to baseline values (p=0.78). Conversely, RSFC between contralesional S1FP and remapped S1FP at 4 weeks in the +Stim group was significantly reduced from baseline (p=0.041) and indistinguishable from week 1 (p=0.79) (**Fig. 4B, D**, **S2**). At 4 weeks, significant disruption of intrahemispheric, ipsilesional RSFC between remapped S1FP and motor was also observed in +Stim mice but not +Stim mice (p=0.015) (**Fig. S2B**).

**Figure 4.**
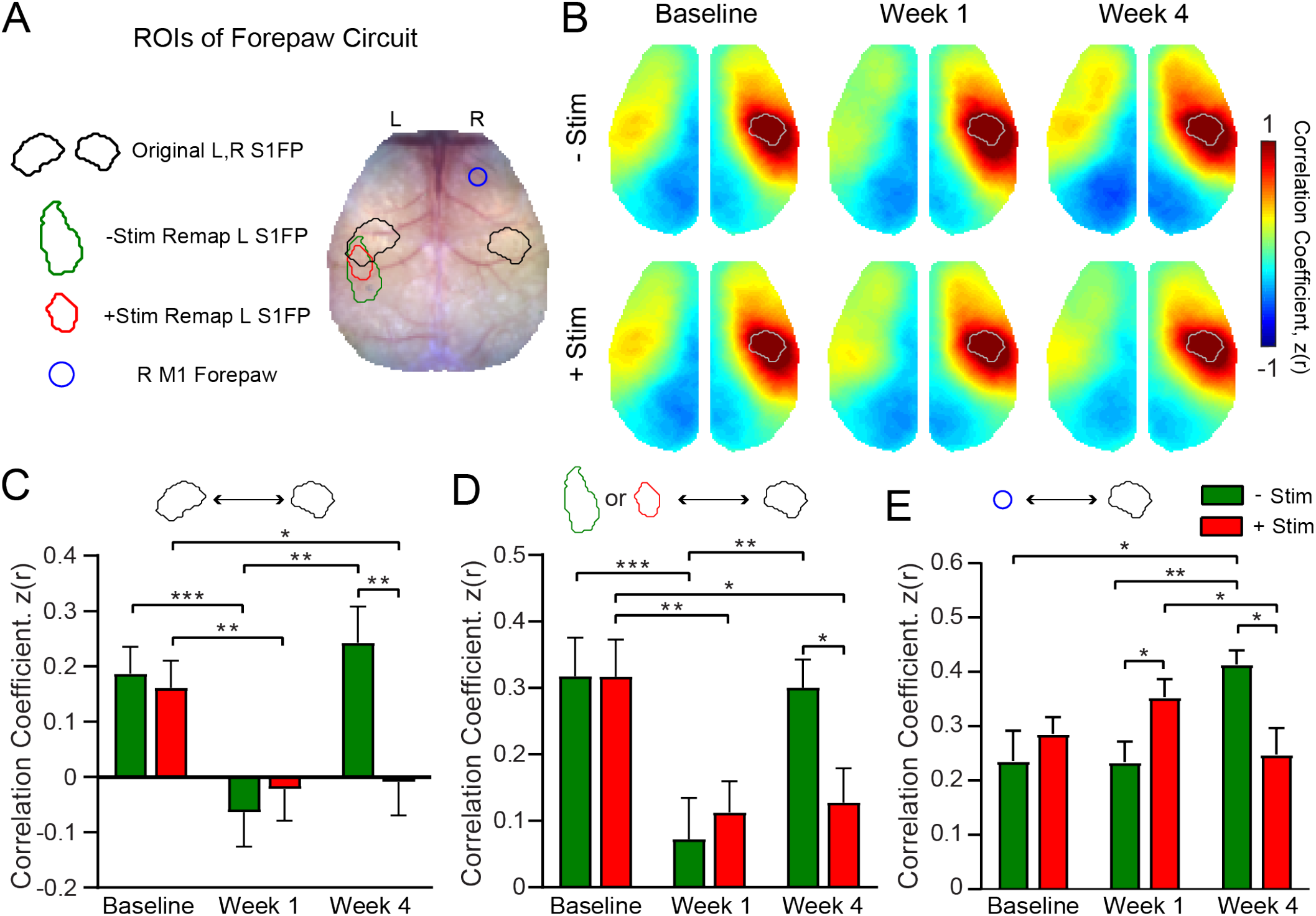
Contralesional stimulation inhibits recovery of somatomotor forepaw circuitry. A) Regions of interest overlaid on a representative white light image of the dorsal mouse skull. ROIs were determined from evoked responses at Baseline (original S1FP in the Left or Right hemisphere) or at 4 weeks after stroke (Remapped Left S1FP). Responses at 4 weeks contained increased activity in Right M1FP and included as an ROI for S1FP RSFC analysis. B) Group averaged maps of RSFC for the Original R S1FP ROI in both –Stim (n=10) and +Stim (n=15) groups reveal strong ipsilateral connectivity with sensory and motor regions, as well as bilateral RSFC with left forepaw cortex. Similar deficits are observed in both groups 1 week after stroke. Bilateral RSFC in both groups is nearly ablated while ipsilateral connectivity remains relatively preserved. By 4 Weeks, RSFC in Right S1FP in the –Stim group exhibits new functional connections in perilesional somatosensory regions, including remapped Left S1FP as well as more anterior motor cortices. Interhemispheric RSFC of Right S1FP in the +Stim group remains largely absent by 4 weeks. C)-E) Quantification of RSFC over time in ROIs depicted in A). C) Homotopic RSFC in original S1FP is significantly reduced in both groups after stoke, and only recovers in –Stim mice: Repeated measures 2-way ANOVA revealed a significant Group x Time interaction (p=0.009) and a main effect of Time (p=0.0003). D) “Homotopic” RSFC with remapped forepaw regions is reinstated by 4 weeks after stroke in –Stim mice only: Repeated measures 2-way ANOVA of revealed a significant Group x Time interaction (p=0.023) and a main effect of Time (p=0.01). E) Ipsilateral RSFC between Right S1FP and Right M1FP. Despite exhibiting increased activity in R right M1FP during left forepaw stimulation, +Stim mice do not exhibit increased ipsilateral RSFC between RM1FP and RS1FP. Repeated measures 2-way ANOVA revealed a significant Group x Time interaction (p=0.0002). All Post hoc tests were performed as t-tests assuming unequal group variance and corrected for multiple comparisons using false discovery rate correction. *=p<0.05; **=p<0.01; ***=p<0.001; ****=p<0.0001.

As reported in Figures 2C, D, electrical stimulation of the unaffected forepaw resulted in a trend towards (−Stim) or significant increase in (+Stim) stimulus evoked activity in contralesional S1FP and M1FP following stroke. We next examined if increased contralesional somatomotor activity was associated with increased intrahemispheric RSFC between these regions (**Fig. 4E**). In +Stim mice, intrahemispheric RSFC between sensory and motor cortices did not appreciably change over time from baseline values, despite exhibiting increased area and magnitude in these regions after photothrombosis. At week 4, −Stim mice exhibited higher intrahemispheric RSFC between sensory and motor regions compared to +Stim mice (p=0.0068) suggesting the involvement of the contralesional hemisphere during spontaneous recovery. At week 1, +Stim mice demonstrated a small but significant (p=0.027) increase in intrahemispheric RSFC compared to –Stim mice. However, this increase did not persist in +Stim mice over time. In control mice, photostimulation did not significantly alter homotopic S1FP RSFC (**Fig. S3A, B**) or intrahemispheric RFC between Right S1FP and Right M1FP (**Fig. S3C**). Together, these results suggest that more complete cortical remapping is associated with a return to more normalized patterns of interhemispheric, homotopic RSFC in the affected circuit. Further, larger patterns of activation in the unaffected hemisphere are not indicative of increased intrahemispheric RSFC within that hemisphere.

### Global network interactions after stroke are suppressed by homotopic contralesional stimulation

Functional network disruption following focal ischemia extends outside of the lesioned territory to brain regions remote from the infarct [31]. Brain networks returning towards more normal patterns of intrinsic organization after stroke (i.e. restored homotopic RS-FC) appear to support better behavioral performance[9, 10]. Because +Stim mice exhibited sustained behavioral deficits and lower S1FP RSFC at week 4 compared to –Stim mice, we hypothesized that contralesional focal excitatory stimulation also altered the degree to which global RSFC renormalized in +Stim mice. Whole cortex correlation matrices were generated for all pairwise comparisons within our field-of-view for both groups at week 4 (**Fig. 5A, B**). Matrices are grouped by functional assignment then organized by hemisphere (left, ipsilesional; right contralesional). Qualitatively, −Stim mice (**Fig. 5A**) exhibit stronger homotopic RSFC (off-diagonal elements) compared to +Stim mice (**Fig. 5B**). Further, ipsilateral anticorrelations between anterior-posterior brain regions are also more pronounced, for example between ipsilesional Motor and Visual regions, in –Stim mice compared to +Stim mice. Group wise correlation differences at week 4 were calculated as –Stim minus +Stim (**Fig. 5C**). Compared to +Stim mice, –Stim mice exhibit increased interhemispheric RSFC between ipsilesional sensory and motor cortex (e.g., Box 1, **Fig. 5C**), homotopic RSFC within the somatosensory and motor networks (e.g., Box 2, **Fig. 5C**), as well as increased intra- and inter-hemispheric RSFC within visual and parietal cortex (e.g., Box 3, **Fig. 5C**). Additionally, mice recovering spontaneously exhibit increased anticorrelations between anterior (e.g. motor and sensory) and posterior (visual) cortices (Box 4, **Fig 5C**).

**Figure 5.**
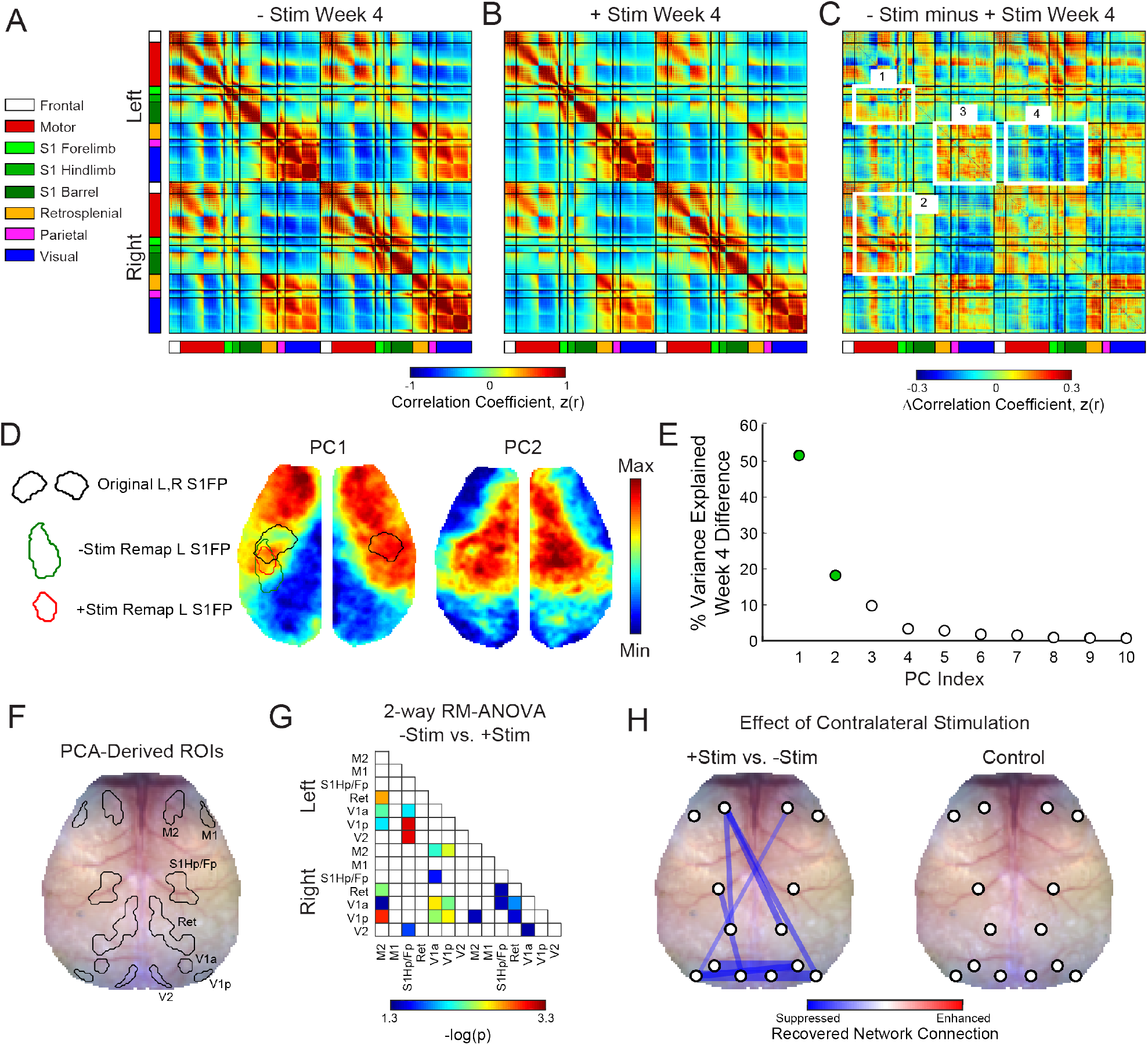
Global network interactions after stroke are suppressed by increased contralesional forepaw activity. Whole cortex correlation matrices for A) –Stim (n=10) and B) +Stim (n=15) groups 4 weeks after photothrombosis. Matrices are grouped by functional assignment then organized by hemisphere (left, ipsilesional; right contralesional). C) RSFC difference matrix (−Stim minus +Stim) shows the group-averaged correlation differences between experimental groups. Notice increased intra- (box 1) and inter-hemispheric (box 2) RSFC within the forepaw somatosensory and motor networks, as well as in distant visual and parietal regions in the −Stim group (box 3). Additionally, mice recovering spontaneously exhibit increased anticorrelations between visual, somatosensory and motor regions (box 4). D) Group-wise RSFC differences examined using unbiased spatial principle component analysis (PCA) of the group-level correlation difference matrix in panel C. First 2 PCs explain 70% of the variance between group differences at 4 weeks. Topography of PC1 reveals increased RSFC of –Stim mice within motor and sensory regions within both hemispheres with a notable island of increased connectivity within the remapped forepaw region. Pronounced anticorrelations can also be observed between posterior visual/retrosplenial regions and anterior sensorimotor regions. PC2 shows that surrounding sensory, medial motor, cingulate and more distant parietal regions also exhibit increased RSFC in –Stim mice compared to +Stim mice. Contours of original and remapped forepaw regions shown for reference. E) The resulting eigenspectrum after PCA of the Week 4 correlation difference matrix in C). PC1 at 4 weeks (green dot) explains 52% of the variance between groups at 4 weeks. F) PCA-derived ROIs for unbiased RSFC analysis. See Fig. S4 for ROI locations in each PC map. G) Repeated measures 2-way ANOVA shows brain wide differences in regional RSFC over time between groups. Regions having a significant Group x Time interaction (p<0.05 uncorrected for multiple comparisons) are shown. Notably, regions outside of the focal injury demonstrate significant differences in RSFC across groups. For example, RSFC between Left M2 and distant retrosplenial and visual regions. Similarly, ipsilateral and bilateral RSFC in Visual and retrosplenial cortices is also significantly altered between groups. H) Left: Stick and ball network diagram showing the effect of stimulation on recovery. Recovery for both groups was defined as the change in magnitude of ROI-based RSFC from Week 1 to Week 4 (see Fig. S4). The difference between these matrices (+Stim minus –Stim) determined the effect of contralesional excitation on spontaneous recovery. Significantly suppressed recovery was observed between nodes connected to the lesion and other regions (left S1hp/fp and left visual; left M2 and left retrosplenial), between nodes distant to the site of direct injury (left and right visual cortex), and across hemispheres (right Motor and left visual). Thicker lines indicate larger changes. Significant differences were those in panel having a significant Group x Time interaction as determined in panel G but corrected for multiple comparisons. Right: RSFC changes in control mice (n=9) over the experiment. No significant changes were observed.

Maps of the most salient group differences in RSFC were created using unbiased spatial principal component analysis (PCA) of the group-level, week 4 correlation difference matrix (**Fig. 5D, E**). PC1 reveals the topography associated with the largest RSFC differences between groups. This map, which explains 52% of the inter-group variance at week 4, includes large portions of motor and somatosensory cortex (reds) and an island of connectivity within the remapped forepaw regions. Pronounced anticorrelations (blues) can also be observed between posterior visual/retrosplenial regions and anterior sensorimotor regions. PC2, which explains 18% of the variance between groups at week 4, shows that perilesional and contralesional sensory, medial motor, cingulate and more distant parietal regions also exhibit increased RSFC in –Stim mice compared to +Stim mice. The resulting eigenspectrum after PCA of the Week 4 correlation difference matrix reveals that, in total, these observations explain 70% of the variance between groups (PC1: 52%; PC2: 18%), **Fig. 5E**).

In order to generate unbiased ROIs for probing evolving global RSFC after stroke, the first 2 PCs shown in Fig. 5D were each averaged across midline, smoothed, and thresholded at a confidence interval of 85% for both positive and negative values. This procedure produced 7 PCA-derived ROIs in each hemisphere corresponding to regions overlapping with primary and secondary motor (M1, M2), primary sensory hindpaw/forepaw (S1hp/fp), retrosplenial (Ret), anterior and posterior primary visual (V1a, V1p) and secondary visual (V2) cortex (**Figs. 5F, S4A**). For each mouse and group, pixel time traces were averaged within each ROI and correlated to generate regional RSFC matrices at each time point for all pair wise comparisons(**Fig. S4B**). Repeated measures 2-way ANOVA applied to each matrix element over time shows brain-wide differences in regional RSFC over time between groups (**Fig. 5G**). P-values are shown for all elements having a significant Group x Time interaction (uncorrected p<0.05). Regions closest to direct injury (i.e., S1HP/FP) exhibited significant differences in RSFC with visual brain regions. Further, RSFC within and across regions distant to the lesion was also significantly affected, for example, inter and intra hemispheric RSFC within visual and retrosplenial cortices, as well as RSFC between visual and motor regions across hemispheres.

Global network recovery was determined in each group by subtracting the magnitude of ROI-based RSFC at week 1 from those at week 4 (**Fig. S4B, C**). Significant differences (FDR corrected p<0.05) were visualized as a stick and ball network diagram on the mouse skull with line thickness indicating connection strength (**Fig. 5H**, left). Comparing recovery matrices of –Stim mice versus +Stim mice revealed that chronic excitation after stroke predominantly suppresses restoration of global RSFC (blue lines, **Fig. 5H**, left). Specifically, +Stim mice exhibited significantly suppressed recovery between nodes connected to the lesion and other regions (left S1hp/fp and left visual; left M2 and left retrosplenial), homotopic nodes distant to the site of direct injury (e.g., left and right visual cortex), and across hemispheres (e.g., right Motor and left visual). The effects of chronic stimulation on evolving global RSFC appear to depend on the post stroke environment. In control mice, chronic Right S1FP stimulation did not significantly alter RSFC between any ROI pairs over the course of 4 weeks (**Fig. 5H**, right).

### Recovery of functional connection density is suppressed by focal contralesional activity

In addition to mapping pairwise RSFC differences across groups, we also examined the number of functional connections (node degree) exhibited by each pixel over the brain (**Fig. 6**) Maps of global node degree for each group and time point were calculated as the number of functional connections exhibited by each pixel having a correlation coefficient z(r)>0.4. At Baseline, both groups exhibit high node degree (reds) in motor, anterior somatosensory, medial parietal, retrosplenial and visual regions, with fewer connections (blues) exhibited by lateral sensory areas (**Fig. 6A**). Focal ischemia in left S1FP reduced node degree in local forepaw circuits, as well as lateral sensory and motor areas with only minimal differences evident across groups at week 1 (**Fig. 6B**). However, the effects of chronic stimulation became profound by Week 4. Substantial increases in node degree are evident in –Stim mice in several brain regions while +Stim mice exhibit topographically similar maps of node degree to those at Week 1. The largest differences in global connection number between –Stim and +Stim mice at week 4 were observed within right forepaw cortex, large portions of motor cortex as well as posterior cingulate, medal parietal, retrosplenial, and visual areas (**Fig. 6C**, week 4). Many of these regions exhibited a significant Group x Time interaction following a 2-way RM ANOVA over the experiment (**Fig. S5A**).

**Figure 6.**
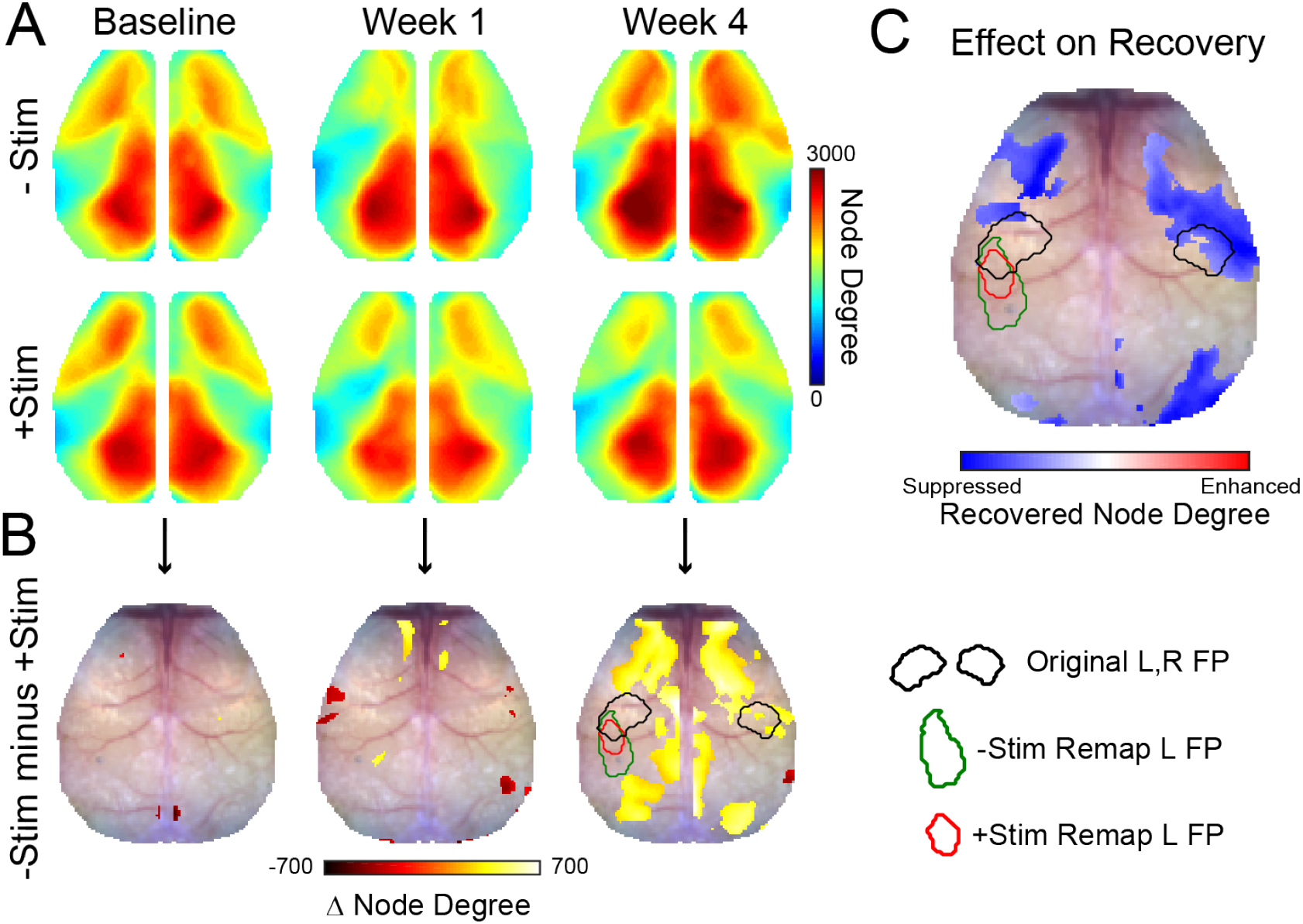

To determine how chronic stimulation altered recovery of regional connection density, maps of node degree at week 1 were subtracted from those at week 4 in each group (**Fig. S5 B, C**). In –Stim mice, the number of functional connections increases over most of the cortex from week 1 to 4 (**Fig. S5B**), whereas in +Stim mice, only portions of parietal and lateral somatosensory cortex exhibit any change. Group-wise differences in recovery (**Fig. 6C, Fig. S5D**) reveal that chronic excitatory activity profoundly suppresses reestablishment of local and global connections within ipsilesional motor, small portions of visual and in the contralesional hemisphere, motor cortex, primary somatosensory forepaw and surrounding cortex, and visual cortices (**Fig. 6C**).

### Contralesional activity inhibits expression of genes important for plasticity after stroke

The lack of local and global reconnection within the somatomotor network of +Stim mice suggests that contralesional stimulation might be affecting molecular mechanisms responsible for the formation of new connections after stroke. Using RT-PCR, we examined gene expression profiles of specific transcripts associated with inflammation, growth, inhibitory/excitatory signaling, synaptic plasticity, dendritic branching, and extracellular matrix remodeling following focal ischemia (**Fig. 7**). Unsupervised hierarchical clustering was used to qualitatively examine broad changes in genetic expression in perilesional and contralesional tissue in both groups (**Fig. 7A**). We identified three larger dendrograms (color coded orange, green, and black) segregating groups of genes based on patterns of expression, and independent of group or tissue assignment. Levels of expression in –Stim mice in either hemisphere provide a “spontaneous recovery phenotype” (−Stim columns, **Fig. 7A**) for evaluating against the effects of contralesional stimulation. The most pronounced differences in genetic expression between groups occurs within the green dendrogram. For example, in ipsilesional tissue, 27 genes upregulated (reds) in –Stim mice were all downregulated (blues) in +Stim mice (green tree, **Fig. 7A**). Further, in contralesional tissue, a subset of these genes were also upregulated in –Stim mice and suppressed in +Stim mice (i.e., 9 rows from CACNG2 through GAD1 within the green tree). Clustering among the 4 groups confirmed that –Stim and +Stim mice have the most disparate expression patterns within ipsilesional tissue.

**Figure 7.**
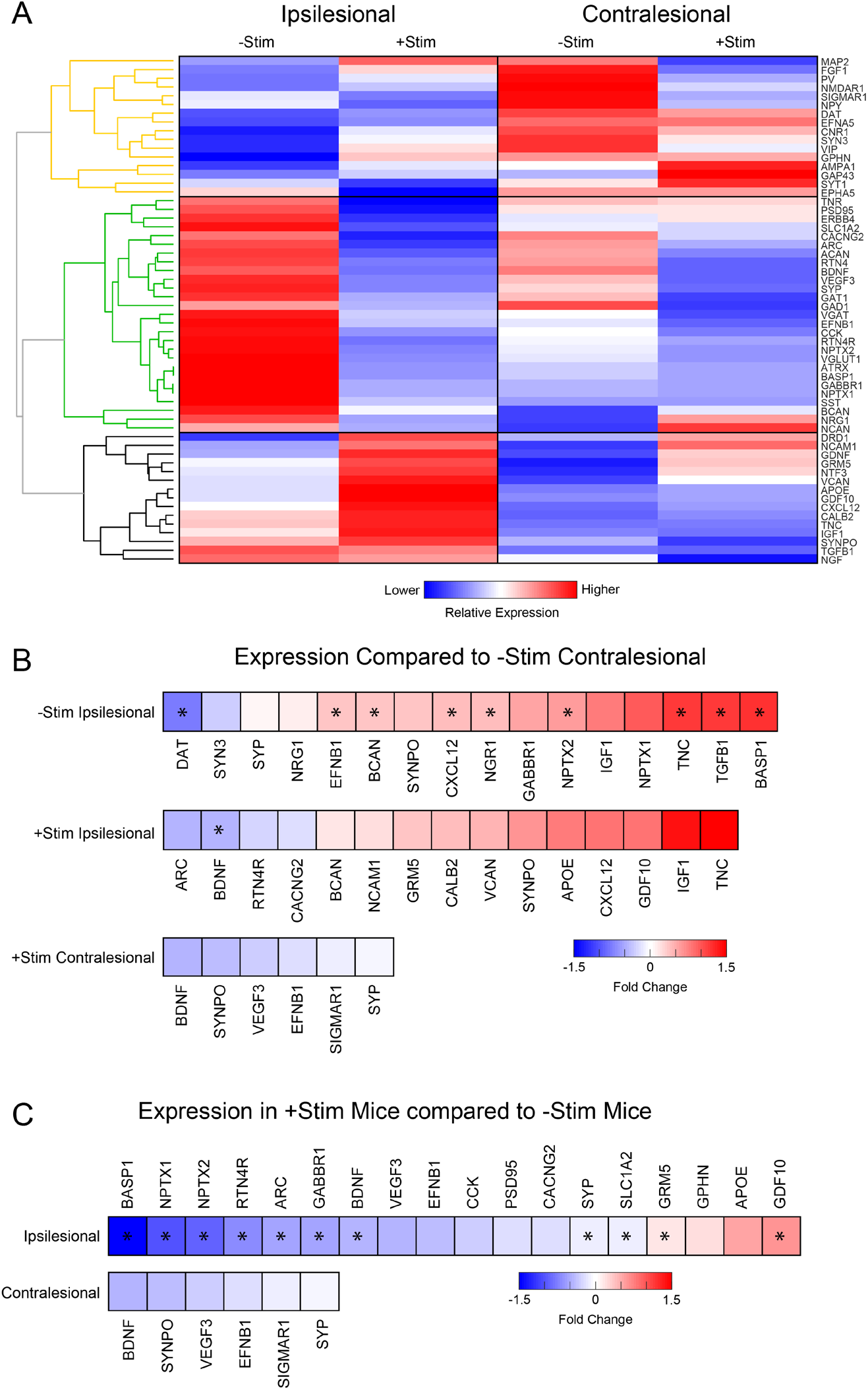
Contralesional activity alters expression of genes important for recovery. Quantitative real time polymerase chain reaction (RT-PCR) analysis was performed in perilesional and contralesional tissue of –Stim (n=5) and +Stim (n=5) mice. A) Broad differences in genetic expression profiles were qualitatively examined with hierarchical clustering. Three larger dendrograms (color coded orange, green, and black) segregate groups of genes based on patterns of expression, independent of group or tissue assignment. Levels of expression in –Stim mice in either hemisphere provide a “spontaneous recovery phenotype”. Within ipsilesional tissue, 27 genes upregulated (reds) in –Stim mice were all downregulated (blues) in +Stim mice (green tree). Further, in contralesional tissue, a subset of these genes were also upregulated in –Stim mice and suppressed in +Stim mice (i.e., 9 rows from CACNG2 through GAD1 within the green tree). Mouse GAPDH was used as a normalization reference. B) Fold change of expression in both groups normalized to expression levels in the contralesional hemisphere of –Stim mice. This analyses allows for comparing expression associated with spontaneous recovery to the combined effect of stroke and stimulation. C) Fold change of expression in +Stim mice compared to –Stim mice in either perilesional or contralesional tissue. This analysis allows for examining the effects of contralesional stimulation after stroke. In panels B and C, colored boxes report fold change of any gene upregulated (reds) or downregulated (blues) with respect to –Stim mice with p ≤0.05 (uncorrected). Starred boxes indicate significance following correction for multiple comparisons (p<0.1 following FDR correction). Contralesional stimulation significantly altered canonical pathways associated with plasticity after stroke.

Two strategies were implemented in order to quantify genetic expression differences across groups. The first method anchored all expression data to the contralesional hemisphere in –Stim mice (**Fig. 7B**). This analyses allows for comparing expression associated with spontaneous recovery to the combined effect of stroke and stimulation. To isolate the effects of contralesional stimulation after stroke, we also compared expression levels in +Stim mice to those in –Stim mice in either hemisphere (**Fig. 7C**). In both analyses, colored boxes report fold change of any gene upregulated (reds) or downregulated (blues) with respect to –Stim mice with p ≤0.05 (uncorrected). Starred boxes indicate significance following correction for multiple comparisons (p<0.1 following FDR correction). Contralesional stimulation significantly altered canonical pathways associated with plasticity after stroke. In perilesional tissue (top row, **Fig. 7C**), this maneuver suppressed genes associated broadly with growth (BDNF), axonal guidance and regulation (BASP1, RTN4), synaptic plasticity and regulation (ARC, SYP, NPTX1, NPTX2), and neuronal signaling (GABBR1, SLCA2). Additionally, contralesional stimulation also resulted in significantly increased metabotropic glutamatergic signaling (GRM5) and some axonal sprouting cues (GDF10). Other transcripts broadly associated with inhibition (GPHN, CCK) or excitation (CACNG2, PSD95) were not observed to be significantly different following correction for multiple comparisons.

## Discussion

Most stroke patients show some degree of recovery, suggesting that endogenous mechanisms of repair are important. Thus, it is essential to understand what processes might inhibit or interfere with spontaneous repair mechanisms. The present work examines how contralesional, excitatory activity in homotopic brain regions functionally connected to the site of injury influences cortical reorganization after stroke. We show that functional recovery occurs in tandem with formation of new cortical representations of the affected limb, restoration of RSFC within the affected somatosensory network, and more global renormalization of functional networks outside of the lesioned territory. Focal excitation of contralesional cortex significantly suppressed these processes, and altered transcriptional changes in several genes important for recovery.

### More complete recovery is associated with more normal patterns of activity in the affected circuit

Spontaneous recovery (i.e. in –Stim mice) was associated with new cortical representations of the right (affected) limb having an area of activation and response magnitude nearer to baseline levels (**Fig. 2A, B).**Conversely, perilesional cortex in +Stim mice failed to remodel as completely and exhibited sustained reductions in evoked activity at week 4 (**Fig. 2A, D**). Stimulation of the left (unaffected) limb resulted in the opposite picture. Responses in –Stim mice were indistinguishable from baseline responses while those in +Stim mice exhibited increased activity in Right S1FP and Right M1FP that persisted over the 4 week recovery period (**Fig. 2C, D, Fig. 3B**) consistent with disinhibition from the lesioned hemisphere[32, 33]. The occurrence of new cortical S1FP representations in –Stim mice was therefore associated with more normal patterns of evoked activity in the larger S1FP circuit. Chronic contralesional excitatory stimulation prevented the formation of new cortical representations of the affected limb, and resulted in abnormally large representations of the unaffected limb. In parallel with cortical remapping, we also observed that the presence of new cortical S1FP representations in −Stim mice was associated with restoration of RSFC amongst functionally homotopic brain regions (**Figs. 4D, S2A**). That is, regions of homotopic synchrony with Right S1FP shift from their original territory in Left S1FP to remapped left S1FP representations. New representations also exhibit similar patterns of RSFC with other brain regions formally connected to the lesioned territory (e.g., left motor). Thus, more complete recovery experienced by –Stim mice was also associated with the reestablishment of more synchronous intrinsic activity within the larger somatomotor network.

The occurrence of new cortical S1FP representations along with a restoration of balanced stimulus evoked activity and homotopic RSFC in the S1FP circuit suggests that cortical remapping of disconnections after stroke provides a clue to how local circuits reconnect with brain networks to facilitate behavior. Following focal ischemia, periinfarct regions appear to undergo large-scale changes in neuronal response properties. For example, in both human and animal studies periinfarct regions become more responsive to stimulation of somatomotor regions with which they are not typically associated [4–6, 34, 35]. In rodents periinfarct remodeling correlates temporally with behavioral recovery [4, 5], and behavioral deficits in rats can be reinstated by ablating these remapped regions[36], suggesting that remapped cortex assumes the function of brain regions lost to stroke. Reestablishing normal patterns of stimulus evoked or resting state activity appears essential for regaining function. Brain networks returning towards more normal patterns of intrinsic organization after stroke (i.e. restored homotopic RSFC) also exhibit more normalized patterns of activation [4, 5, 37–40]. Abnormal intrinsic brain rhythms that give rise to abnormally strong patterns of task activation result in reduced variability of motor patterns after stroke[41]. Abnormal brain activity could reflect an attempt to link, albeit inefficiently, regions that are disconnected, either by increasing neural activity upstream of the lesion [28] or by rerouting activity through accessory regions [32, 41]. Intermittent contralesional excitatory stimulation could exacerbate these processes.

### Increased contralesional activity negatively affects global network interactions

Ischemic stroke results in direct structural damage to local brain networks but also to functional damage to global networks outside of the lesioned territory[1, 2, 42]. While brain network dysfunction after stroke appears to be largely due to structural damage [31, 43-46], widespread functional disruptions are topographically linked within functional networks that can span across hemispheres [31]. Because functional outcome after stroke depends on the local site of ischemic injury as well as on remote connections [47–49], It is therefore necessary to consider more than just the ischemic territory when characterizing systems-level mechanisms of brain repair[50]. ROI-based RSFC analysis of the somatomotor network shows clear changes in network synchrony over the 4 week recovery period in –Stim mice (top row, **Fig. 4B**) that were not observed in +Stim mice (bottom row, **Fig. 4B**). Beyond increased RSFC between Right and remapped Left S1FP at week 4, Right S1FP exhibits higher RSFC with Right M1FP and with regions anterior to the infarct in motor cortex. Supporting this qualitative observation, maps of node degree in –Stim mice indicate that large portions of motor and sensory cortices exhibit more functional connections than +Stim mice (**Fig. 6**). Additionally, regions well outside of direct injury (e.g. Visual, Retrosplenial) also exhibit changes in functional connection number. These data suggest that both local and distant networks are altered after stroke, and that the degree to which these networks recover is impaired by increased contralesional stimulation (**Fig. 6C**). However, these findings do not offer insights into the effect of increased contralesional activity on evolving internetwork interactions during recovery. To characterize global network interactions, we examined all pairwise RSFC over the cortex (**Fig. 5**). Unexpectedly, several brain regions underwent dramatic changes in RSFC structure. In –Stim mice, Parietal, Visual and Retrosplenial cortices all exhibited increased homotopic RSFC (**Figs. 5C, D, G**) compared to +Stim mice. Further, +Stim mice exhibited significant reductions in the magnitude of anticorrelations between functionally opposed networks residing within anterior (Primary and secondary motor) and posterior (Retrosplenial, Primary visual) cortices (**Figs. 5B, D, G**). Dynamic patterns of anticorrelated activity segregate distinct processing streams across networks during both task and resting state paradigms[51, 52]. In humans, the most prominent example occurs between the dorsal attention network and default mode network. Altered synchrony across large cortical distances (>several mm), as observed by reduced anticorrelated activity in +Stim mice at Week 4, could prevent inter-network communication necessary for higher-order integration. Reduced node degree in +Stim mice might indicate network-level inefficiencies important for segregation, exemplified by more asynchronous intrinsic activity within these networks. We have recently shown[53] that improved tactile proprioception after stroke was associated with increased node degree in motor, somatosensory, and parietal cortices in mice, all regions relevant for processing proprioception and touch[54]. Reduced within-network connectivity (i.e., global node degree) combined with poorer interactions across networks (i.e. reduced anticorrelations) could impair multi-domain behavioral output in mice after stroke.

The findings that reconnection between remodeled cortex and homotopic brain regions parallels global network renormalization implies that these processes are both required for proper recovery. Further, the fact that focal excitation of contralesional excitatory circuits significantly suppresses local and distant network communication suggests that even small changes in contralesional activity after stroke can significantly impact recovery trajectories. Recent mouse work probing causal relationships between local neural stimulation and recovery of specific (sensorimotor) behaviors further support the idea that more normal patterns of activity are associated with better behavior. For example, photostimulation of the ipsilateral primary motor cortex in transgenic Thy1-ChR2 mice promoted functional recovery and a return to normal patterns of optogenetically-evoked perfusion changes in perilesional cortex [25]. In a separate study, optogenetic stimulation of thalamocortical axons also promoted the formation of new and stable thalamocortical synaptic boutons, which was correlated with enhanced recovery of somatosensory cortical circuit activity and forepaw sensorimotor performance [28].

### Contralesional stimulation alters transcriptional changes in genes important for recovery

Recovery from stroke is associated with neuronal and synaptic plasticity, the formation of new structural and functional connections that take over those lost due to infarction. These changes are likely to occur both locally (at the peri-infarct region) and at distant sites where changes in connectivity may occur (e.g. contralesional hemisphere). We examined changes in the expression of select genes associated with neuronal plasticity to determine if contralesional stimulation altered expression of genes associated with spontaneous recovery. While distinct phases of plasticity occur over different time scales[55], good recovery of the periinfarct is associated with upregulation of neurotrophic signaling[56], growth-associated gene expression[57], and markers of axonal formation/migration[58] and synaptogenesis[1, 12]. Some of these repair-related molecular changes are also observed in homologous, contralesional sites[3].

Our cumulative results suggest that spontaneous recovery following focal ischemia may be suppressed by chronic contralesional excitation. We used cluster analysis to determine if changes in neuroplasticity genes associated with spontaneous recovery (−Stim group) were suppressed by chronic contralesional excitation. Four groups were entered into the cluster analysis (ipsilesional and contralesional tissue in both groups). Hierarchical clustering revealed three distinct clusters that segregated patterns of gene expression. Of particular interest were a collection of genes that demonstrated increased expression in ipsilesional cortical tissue in the –Stim group, but were suppressed in the +Stim group: BASP1, NPTX1, NPTX2, ARC, BDNF, SYP, and SLC1A2. Interestingly, a subset of these genes were also increased in the contralesional tissue, and also suppressed in the +Stim group (ARC, BDNF, SYP, SLC1A2). In line with the findings, we recently showed that remapping of forepaw cortex is dependent on activity-dependent synaptogenesis, and that blocking ARC-dependent remapping limited behavioral recovery[12]. NPTX1 and 2 mediate synaptic clustering of AMPA receptors, are implicated in synapse formation and maintenance [59, 60] and facilitate metabotropic glutamate receptor-mediated long-term depression[61]. Increased excitatory input from the contralesional hemisphere chronically after stroke appears to modify these repair-related molecular events.

Altering cell signaling, or modulating inhibitory or excitatory tone can profoundly affect recovery [62–64]. The negative effect of contralesional excitation on recovery is thought to be due to increased perilesional inhibitory influences. While we did survey for markers of inhibition and excitation (**Table S1**), most were not observed to be significantly altered between groups after correcting for multiple comparisons. However, stimulated mice did exhibit significant reduction in ipsilesional expression of GABABR1, a G protein-coupled receptor abundantly localized to glutamatergic synapses which regulates excitatory neurotransmission and synaptic plasticity[65]. Alone, focal ischemia results a loss of GABA B -mediated interhemispheric synaptic inhibition in stroke periphery[64]. Sustained activation of glutamate receptors following focal ischemia leads to rapid downregulation of GABA B receptors via lysosomal degradation[66, 67]. Glutamate-induced down-regulation of GABABR1 would lead to reduced inhibition of glutamatergic synapses, increasing synaptic excitability potentially to the point of increased excitotoxicity[66]. While we did not observe group-wise differences in lesion volume, future studies are required to determine whether increasing ipsilesional inhibition during the recovery phase[68, 69] offsets the effects of contralesional excitation.

Other genes that have been reported to be important for post-stroke plasticity and recovery were either not included in this cluster or exhibited changes in expression that are more difficult to interpret. For example, in the post-stroke axonal sprouting transcriptome, GDF10 is one of the most highly upregulated genes during the initiation of axonal sprouting in peri-infarct cortical neurons[70]. Additionally, RTN4R forms part of a signaling complex (i.e. Nogo) that inhibits axonal growth and limits experience-dependent neural plasticity[71]. Despite a lack of remapping in +Stim mice, this group exhibited significantly increased expression of GDF10 and decreased expression of RTN4R in perilesional tissue. Alone, these changes might facilitate perilesional remodeling. However, new axonal growth also requires other markers that were either unchanged (e.g. GAP-43[57]) or significantly reduced (BASP1[72]) in this group. New axonal growth and subsequent navigation likely require coordination across of several molecular cues to ensure proper synaptic targeting. It is also important to note, that the current analysis was performed at a single time-point (14 days after stroke), during dynamically changing molecular environment[55].

### Interhemispheric influences on recovery and implications to clinical stroke

In one model of information exchange across hemispheres, excitatory information from one hemisphere conveyed by the corpus callosum activates inhibitory cells in the opposite hemisphere, thus inhibiting activity[73, 74]. This inhibition between the left and right hemispheres may be important for bilateral dexterity in upper and lower limb movements, and is required for integrating bilateral sensory and motor signals[75, 76]. Unilateral injury (e.g. stroke) modifies interhemispheric connectivity in such a way that disrupts the balance across hemispheres by decreasing the inhibitory influence of the affected hemisphere [33, 77, 78]. Non-invasive brain stimulation techniques, such as rTMS and tDCS, have been used to influence rehabilitation after stroke by affecting this E/I balance. However, evidence supporting the clinical use of these techniques has been inconsistent. Some clinical trials have shown that these methods have a large, positive effect on motor recovery[21], while other studies were inconclusive[22]. A portion of the variability in outcome is due to grouping patient populations with different types of strokes (ischemic vs. hemorrhagic) and/or with lesions in different locations (cortical vs. subcortical). Further, the lack of refinement for spatial and cellular specificity, limits the use of rTMS and tDCS for examining specific populations of neurons or connections important for recovery.

Utilizing optogenetic targeting, we examine the role of contralesional, homotopic excitation on functional brain organization after stroke, a maneuver mimicking the chronic use of the unaffected limb. Sustained, increased activation of contralesional M1 using rTMS or through the use of the “good limb” is associated with poorer clinical outcome [14, 15], findings corroborated in rodent models[79–81]. We establish that the observations of poorer outcome following this maneuver are associated with a lack of reconnection of both local circuits lost due to stroke, and the subsequent reintegration of these circuits into global resting state networks. However, the role of the ipsilesional vs. contralesional hemispheres on stroke recovery depends on the size and location of the infarct (for review see [82]) and timing of rehabilitation can profoundly alter outcome[40, 83]. Because structural and functional asymmetries persist in well-recovered stroke patients even after clinical symptoms have normalized[84], more systematic approaches are required to unravel causal influences for differences in outcome.

### Limitations and Future Work

The design of the present study was intended to examine how early, chronic stimulation affected cortical remodeling after stroke. The duration of the experiment did not include time points when spontaneously recovering mice have been shown to exhibit more complete recovery (e.g out to 8 weeks or longer[4]). Our assessments of cortical remodeling and network reorganization were based on systems-level, blood-based measures of brain activity. Visualizing more direct measures of neural activity, e.g. through the use of genetically encoded calcium indicators[85, 86] or voltage sensitive dyes[32], would allow for examining evolving neural network interactions as they relate to behavior[87], or evolving microvascular function[88, 89]. Previous studies have shown that the laser light alone can affect cerebral flow in naive animals [90]. While we did not specifically test if the photostimulation design affected blood flow in the mice used in this study, it did not affect measures of RSFC in controls. Further, previous reports by our lab have shown that similar photostimulus parameters do not affect hemodynamic measures of brain activity in several transgenic mouse strains expressing ChR2 in different neural populations [91, 92]. Finally, the limited scope of our RT-PCR analysis provided associations between expression of select neuroplasticity genes and systems level changes across groups post stroke. More complete evaluations of the post stroke transcriptome, and conformation as to which genes and proteins are upregulated following contralesional stimulation will allow for a more complete understanding of how this maneuver affects cortical remodeling after stroke, or how stimulation alone affects gene expression.

## Materials and methods

### Mice

A total of 89 adult male mice expressing channelrhodopsin (ChR2) under the mouse calcium/calmodulin-dependent protein kinase II alpha (CamK2a) promoter (CamK2a-ChR2) were used for experimentation. CamK2a-ChR2 mice were generated using Cre-Lox recombination (Parent strains: CamK2a-Cre, B6.Cg-Tg(Camk2a-cre)T29-1Stl/J, Stock number: 005359; Lox-ChR2, B6;129S-Gt(ROSA)26Sortm32(CAG-COP4*H134R/EYFP)Hze/J, Stock number: 012569, The Jackson Laboratory). Mice were housed in enriched environment cages and given food and water *ad libitum* with a 12hr On:12hr Off light cycle. 80 mice were used for stroke recovery experiments, and 9 mice were used as controls. 10 mice were euthanized 2 days after photothrombosis for infarct volume quantification. 10 mice (5 −Stim, 5 +Stim) were euthanized 14 days after photothrombosis for qPCR. 35 mice (15 –Stim, 20 +Stim) were used for behavioral testing. OISI data were collected on the remaining mice (9 controls, 10 −Stim, 15 +Stim) (see below and **Figure 1**). All experimental protocols were in compliance with the Institutional Animal Care and Use Committee at Washington University in St. Louis.

### Enriched Environment

Enriched housing has been shown to improve functional recovery after stroke[93, 94], thus increasing the dynamic range of assessments of functional recovery. Mice were housed in 24” × 17” × 8” cages (Nalgene) for 1 month prior to and during the entirety of the experiment. Cages contained: Nestlets, Mouse Arch, Mouse Huts, Mouse Igloos, Mouse Tunnels, Fast Trac (bio-serv) and Enviro-dri crinkle paper. These components were re-arranged or replaced every 1-2 weeks to provide new stimuli for the mice.

### Animal Surgery

Prior to imaging, a clear Plexiglas window was placed on the exposed skull as per our previous studies [95]. A single injection of buprenorphine-SR (1.0 mg/kg SC) was given one hour prior to the surgery. Mice were anesthetized with 3% isoflurane inhalation in air for induction, and kept at 1.5% for the duration of the surgery. A clear Plexiglas window with predrilled holes, sterilized with Chlorhexidine and rinsed with sterile saline, was secured to the scalp with dental cement (C&B-Metabond, Parkell Inc., New York, USA). Mice were monitored for 2 days post-surgery. No imaging was performed during this post-operative period.

### Photothrombosis

Focal ischemia was induced via photothrombosis as previously described [12]. Under isoflurane anesthesia (3.0% induction, 1.5% maintenance), mice were placed in a stereotactic frame. A 532nm green DPSS laser (Shanghai Laser & Optics Century) collimated to a 0.5mm spot was centered on left S1FP (0.5mm anterior to bregma, 2.2mm left of bregma) at low power (<0.25mW). The laser was turned off, and mice were then given 200uL Rose Bengal dissolved in saline (10g/L) via I.P. injection. After 4 minutes, the laser power was set to 23mW and illuminated left S1FP for 10 minutes.

### Optogenetic Photostimulation

A randomized subset of mice was subjected to chronic, intermittent optogenetic photostimulation of homotopic S1FP in the right hemisphere for 5 consecutive days/week for 4 weeks beginning 1 day after photothrombosis. Photostimuli consisted of 20ms pulses delivered at 10Hz for 1 minute on, 3 min off, 1 min on, 3 min off, 1 min on (3 minutes/day) as per a previous study reporting functional improvement when this stimulus was delivered to perilesional tissue [25]. The laser power ranged between 0.2mW - 1mW and was set to a level just below that which elicited overt behavioral output (e.g. forepaw or whisker motor movements in sync with stimuli).

### Electrical Forepaw Stim

Transcutaneous electrical stimulation was applied to the left and right forepaws as previously described [12] by placing microvacular clips (Roboz) on either side of the wrists. Electrical stimulation was provided in a block design (AM Systems Model 2100) with the following parameters: 5 seconds rest, 10 second stimulation (0.5 mA, 0.3ms duration, 3Hz) followed by 35 seconds of rest as previously described [13]. Fifteen minutes of data were collected for each paw (18 trials per paw total).

### Optical Intrinsic Signal Imaging (OISI)

Mice were anesthetized by intraperitoneal injection (i.p.) of a ketamine/xylazine cocktail (86.9mg/kg ketamine, 13.4mg/kg xylazine). To facilitate longer imaging times, after the initial bolus, mice were infused subcutaneously with a saline-ketamine cocktail (34.8 mg/kg/hr ketamine) during the imaging sessions as previously described[13]. A heating pad kept at 37°C maintained the mouse body temperature. Sequential illumination was provided by a custom ring of LEDs centered at four wavelengths (478nm, 588nm, 610nm, and 625nm). A cooled, frame-transfer EMCCD camera (iXon 897, Andor Technologies) captured diffuse light reflectance from the skull over a field-of-view of approximately 1 cm^2^. Data were binned 4×4 on camera, resulting in a frame rate of 120 Hz (Hemodynamic imaging at 30Hz). Mice were imaged before, and 1 and 4 weeks after photothrombosis. Thirty minutes of activation data (15 min per paw, 18 stimulus presentations per paw) and up to 45 minutes of resting state data were collected for each mouse in 5-minute data sets (75 min of data total per mouse).

### Image Processing

#### Initial data reduction

Data from all mice were subject to an initial quality check prior to spectroscopic analysis as per our previous reports[95, 96]. Raw reflectance was converted to changes in hemoglobin (Hb) concentration at each pixel and each time point as we have previously reported[95, 96]. Each pixel’s time series was downsampled from 30 Hz to 1 Hz, and data were filtered between 0.009Hz and 0.08Hz for RSFC analysis and 0.009Hz-0.5Hz for task based measures. Global signal regression was performed prior to any analysis. All imaging data were affine-transformed to a common atlas space determined by the positions of the junction between the coronal vessel separating the olfactory bulb from the cortex and sagittal suture along midline and lambda as we have done previously [95, 96].

#### Stimulus evoked somatosensory forepaw maps

Unless otherwise stated, total hemoglobin was used as contrast because it offers the highest contrast-to-noise[91], is more spatially specific than oxygenated or deoxygenated hemoglobin [41, 91] and is most closely linked to underlying neural activity [97]. For each stimulation block, baseline images (1-5s before stimulation) were averaged together and subtracted from the all images in the block. Stimulation blocks for each paw were averaged together. Maps of peak responses were calculated by averaging images 2 seconds before and 2 seconds after stimulation offset and used for activation area and magnitude calculations. Because evoked responses at early time points post stroke can be difficult to detect, group averaged maps of peak responses at baseline were thresholded at 75% of maximum to define a threshold for what constituted a response for all mice at all time points. The maximum response above this threshold was used for activation magnitude calculations. If no pixels were above this threshold, response area and magnitude were set to zero. Group activation incidence images at each time point were created by binarizing all pixels above the 75% threshold. To calculate the map of Cohen’s D at 4 weeks post stroke, evoked response T-maps were created across stimulus blocks for each mouse. All pixels having a T-value >2 were included.

#### Resting State Functional Connectivity Analysis

Regions of interest (ROIs) for Left and Right forepaw (original or remapped) were determined by group-averaged activation maps from the Baseline and 4-week time points and all pixels within the thresholded activations (see above) were averaged to create time courses for each ROI. Forepaw motor representations were determined by stimulation experiments at week 4. ROI traces were correlated with all other time traces in the shared brain mask to create maps of RSFC for the local forepaw somatosensory circuit. Global RSFC circuits were evaluated via zero-lag correlation for all pixel pairs within the shared brain mask for each mouse. Spatial principal components analysis (PCA) was performed on the group averaged correlation difference matrix at 4 weeks to evaluate the largest sources of variance between the –Stim and +Stim groups as we have done [98]. From this process, the first 2 PCs were used to define ROIs for RSFC analysis. Maps of PC1 and 2 were symmetrized about midline, smoothed and thresholded at 85% of maximum for positive and negative values resulting in 7 ROIs within each hemisphere[98]. Maps of global node degree were calculated as described previously[53, 93] by thresholding whole cortex correlation matrices at z(r) ≥ 0.4. Connections above this threshold were set to 1, and summed over both hemispheres to produce a binarized measure of global node degree for all pixels within the shared brain mask.

### Infarct Quantification

Mice (n=5 –Stim, and n=5 +Stim) were deeply anesthetized with FatalPlus (Vortech Pharmaceuticals, Michigan, USA) and transcardially perfused with heparinized PBS. The brains were removed and fixed in 4% paraformaldehyde for 24 hours and transferred to 30% sucrose in PBS. After brains were saturated, they were snap-frozen on dry ice and coronally sectioned (50 μm thick slices spaced 300 μm apart) on a sliding microtome. After mounting, slices were allowed to dry for 24 hours and stained with cresyl violet, and cover slipped. Stained sections were imaged with a NanoZoomer under bright field setting and infarct area was quantified in ImageJ for each slice. Infarct volume was calculated as total infarct area*300 μm.

### Brain extraction and RT-PCR

Mice (n=5 –Stim, and n=5 +Stim) were deeply anesthetized with isoflurane and transcardially perfused with 0.01M PBS. Periinfarct and contralesional tissues were dissected, snap-frozen on dry ice, and stored at −80°C. Total RNA was extracted using RNeasy Mini Kit (Qiagen) and reverse transcribed with the cDNA Reverse Transcription kit (Invitrogen). Quantitative PCR was performed with SYBR Green using the ABI QuantStudio™ 12K Flex system in the default thermal cycling mode. Mouse GAPDH was used as a normalization reference. Relative mRNA levels were calculated using the comparative Ct method. Prior to clustering, expression levels of each gene were mean subtracted across groups and normalized to unit variance. Genes examined are tabulated in **Table S1**.

### Cylinder Rearing

Cylinder rearing recording and analysis was done as previously described [12, 99]. Briefly, mice receiving photothrombosis were placed in a 1000 mL glass beaker and recorded for 5 minutes. A blinded observer manually analyzed videos to determine the amount of time that the 1) right paw, 2) left paw, or 3) both paws made contact with the glass walls. Paw-use asymmetry was calculated as (% left paw use – % right paw use)/ (% left paw use + % right paw use).

### Statistical Analyses

All analyses were performed in either Graph Pad Prism 8 or Matlab 2018a. All Pearson-R values were converted to Fisher-Z values prior to statistical testing. Differences between stroke groups for imaging measures and behavioral measures were analyzed using 2-way, repeated measures ANOVA (rmANOVA) and corrected for assumptions against symmetry. In the control group, differences across time were assessed using 1-way rmANOVA. Maps of statistical differences were corrected on a clusterwise basis. Significant effects were evaluated post-hoc; these and other pairwise comparisons were performed using a 2-tailed Students t-test assuming unequal group variance and corrected for multiple comparisons by controlling the false discovery rate (FDR). For all tests, significance was achieved if (corrected) p<0.05 except for RT-PCR data (p<0.1).

## Acknowledgements

This work was supported by National Institute of Health grants RO1-NS102870 (AQB), K25-NS083754 (AQB), R37NS110699 (JML), R01NS084028 (JML), R01NS094692 (JML), R01NS078223 (JPC), P01NS080675 (JPC) R01NS099429(JPC), F31NS089135 (AWK) and F31NS103275 (ZPR), the McDonnell Center for Systems Neuroscience (AQB), The Alborada Trust (TW), The Wachtmeister Foundation (TW) and The Swedish Research Council (TW). We also thank Grant Baxter and Jasmine Park for their assistance with imaging data acquisition, and Shannon Macauley for helpful discussions on reporting RT-PCR data.

## SUPPLEMENTAL MATERIAL

**Table S1.**
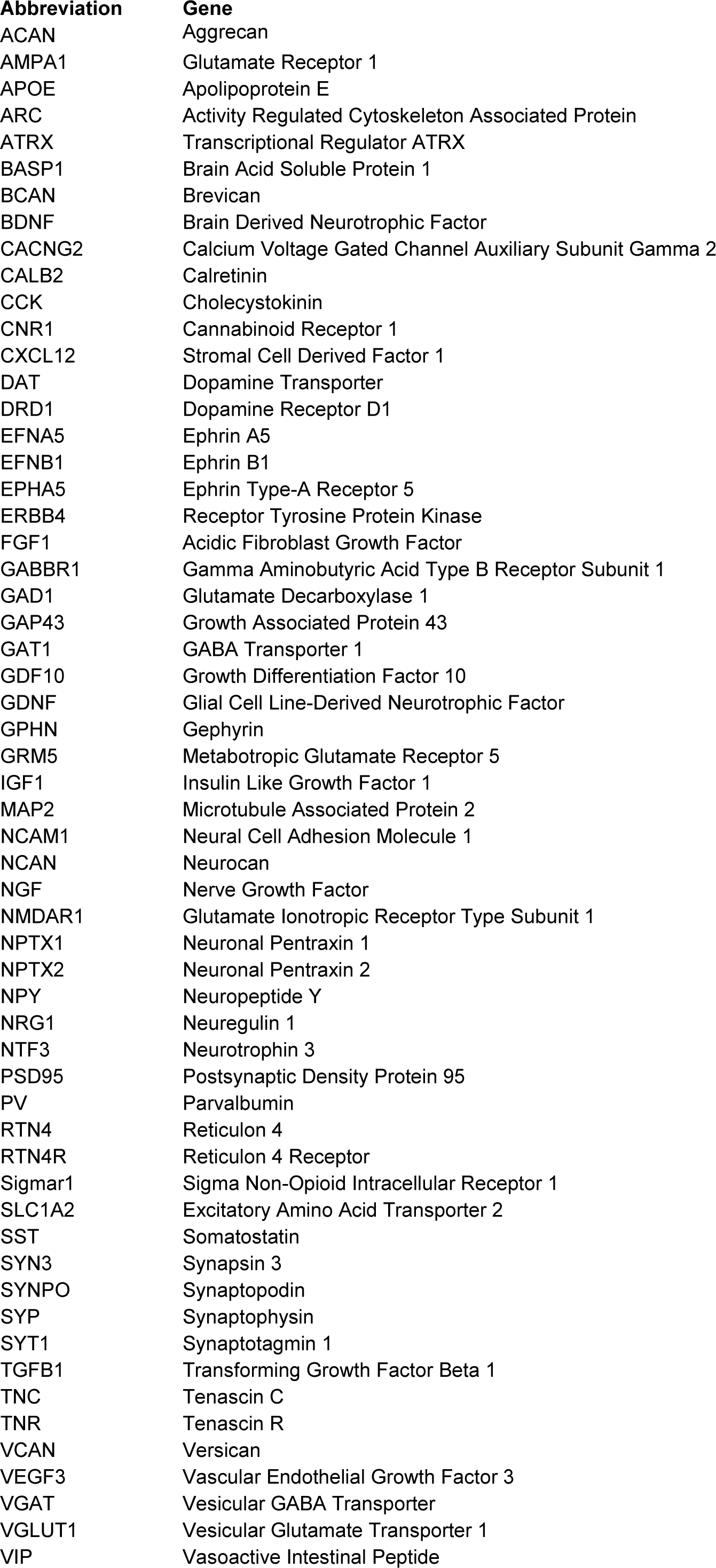
Genes and their abbreviations.

**Figure S1.**
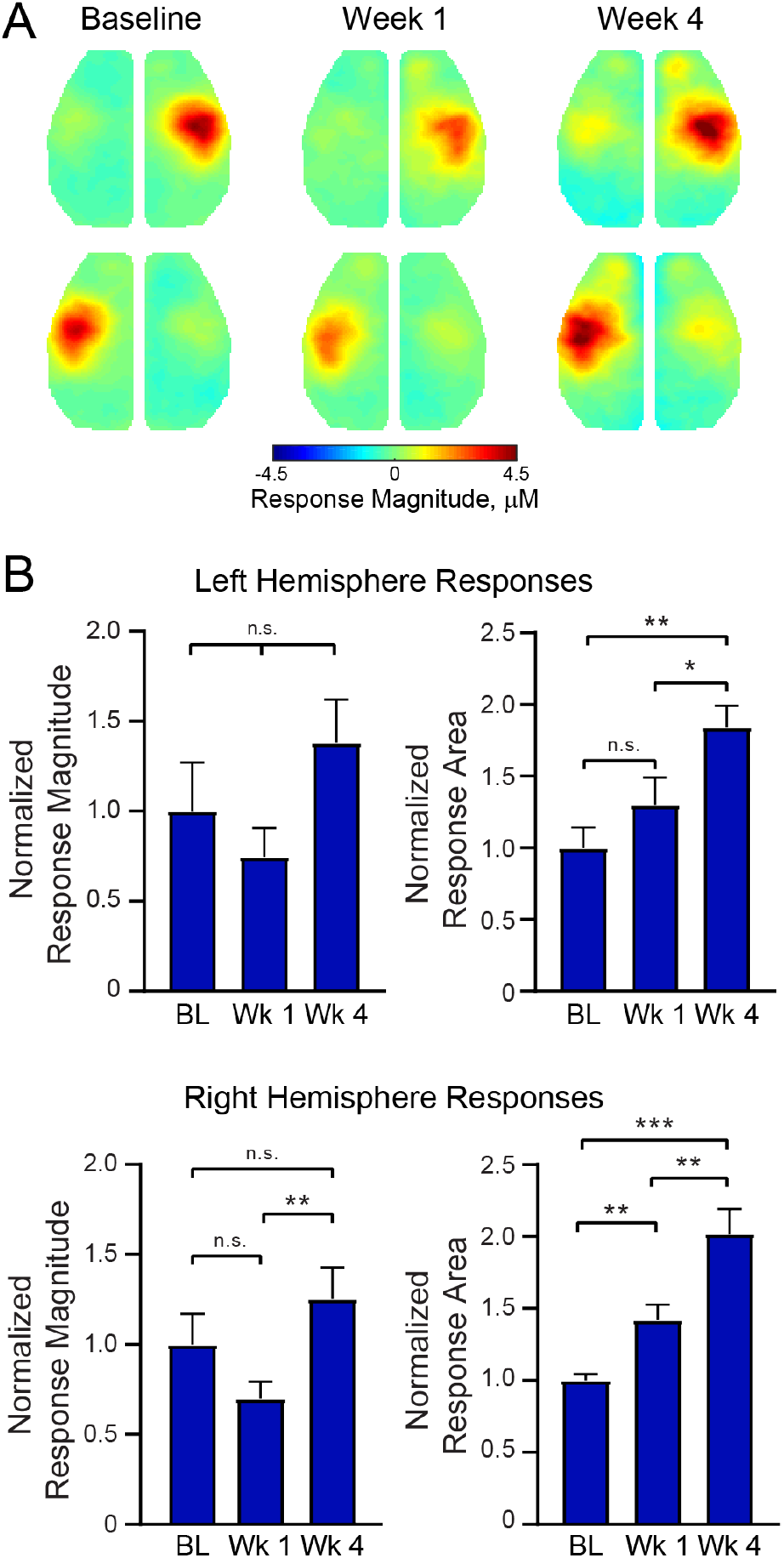
Focal excitatory stimulation influences stimulus evoked responses in healthy mice. A) Cortical responses in mice subjected to the same photostimulation paradigm but without photothrombosis (n=9). Robust responses are observed following stimulation of either limb, and appear to increase in area and magnitude by Week 4 compared to baseline responses. At 4 weeks, note the presence of evoked activity in motor forepaw cortex. B) Quantification of evoked response magnitude and area. Top Row, Left hemisphere: Repeated measures 1-way ANOVA revealed a significant effect of time (p=0.01) for evoked response area. Bottom row, Right hemisphere: Repeated measures 1-way ANOVA revealed a significant effect of time for evoked response area (p<0.0001) and a trend (p=0.063) in evoke response magnitude. Post hoc tests were performed as t-tests assuming unequal group variance and corrected for multiple comparisons using false discovery rate correction. All analysis performed using total hemoglobin as contrast. *=p<0.05; **=p<0.01; ***=p<0.001; ****=p<0.0001.

**Figure S2.**
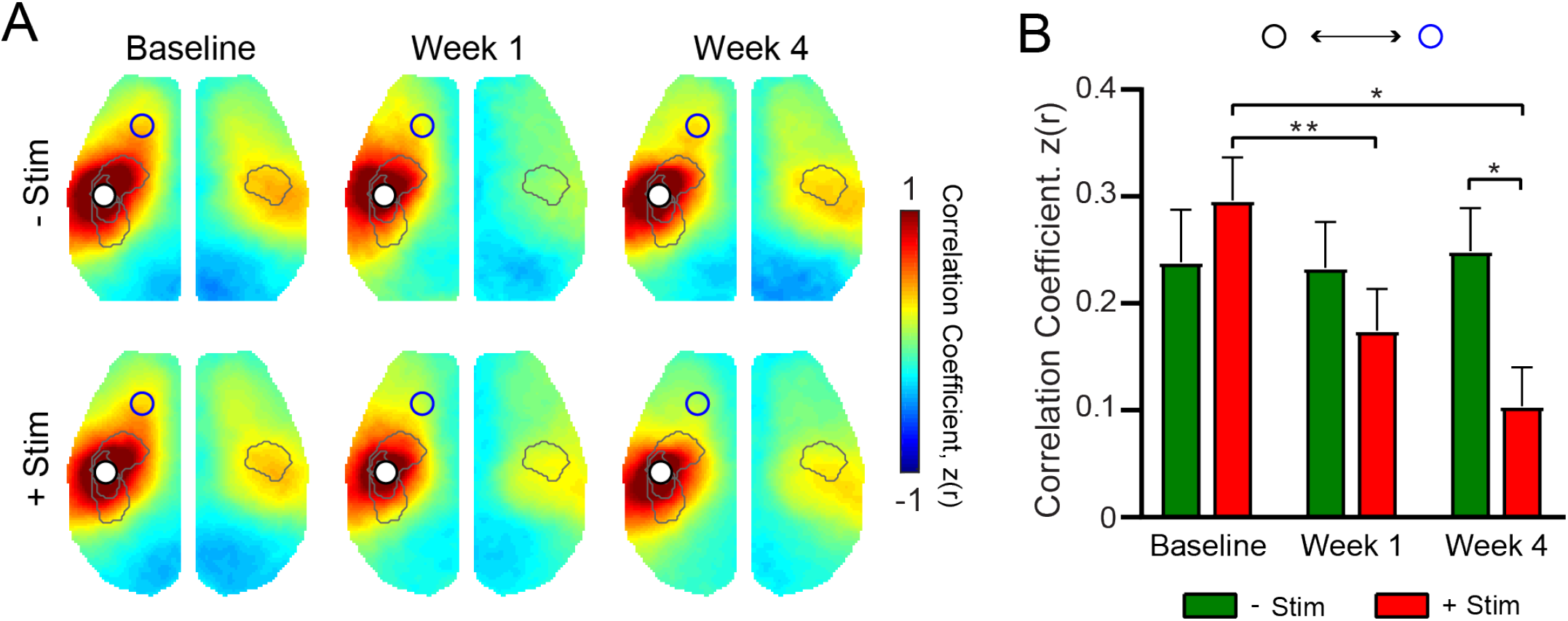
RSFC in Remapped S1FP. A) RSFC maps for remapped S1FP of both groups (−Stim (n=10) and +Stim (n=15)). Cortical region of remapping exhibits strong ipsilateral RSFC and homotopic RSFC with contralateral cortex prior to stroke. At week 1, RSFC between this region and contralateral Right S1FP cortex is significantly disrupted in both groups(See Fig. 4D for quantification). By week 4 robust RSFC with Right S1FP is observed in the –Stim group only. B) Quantification of intrahemispheric, ipsilesional RSFC between remapped S1FP and left motor cortex (similar to panel Fig. 4E). 2-Way, repeated measures ANOVA revealed a significant Group x Time interaction (p=0.014), as well as a significant effect of Time (p=0.028). All Post hoc tests were performed as t-tests assuming unequal group variance and corrected for multiple comparisons using false discovery rate correction. *=p<0.05; **=p<0.01; ***=p<0.001; ****=p<0.0001.

**Figure S3.**
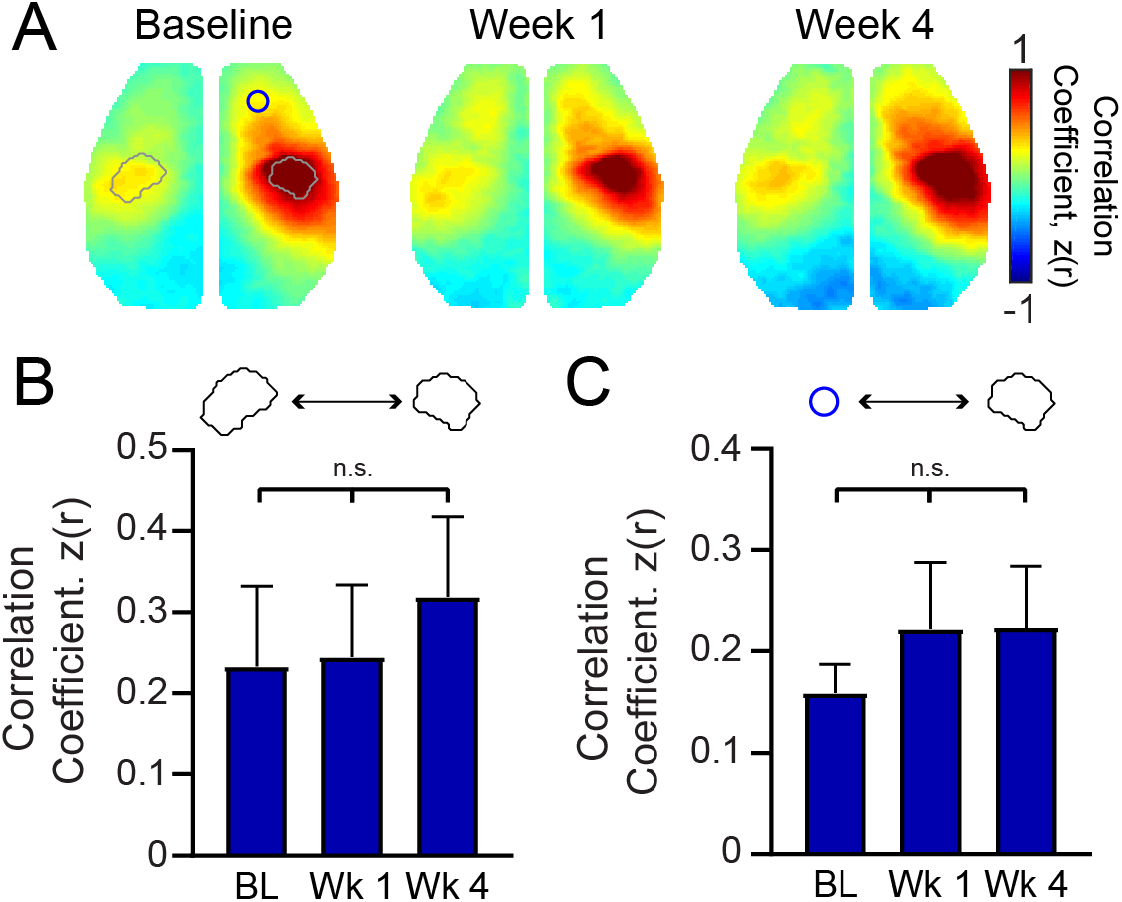
Focal excitatory stimulation in control mice does not affect RSFC in the somatomotor circuit. A) Group averaged patterns of RSFC in control mice (n=9) subjected to the same 4-week long photostimulation paradigm but in the absence of stroke. Regions of interest overlaid on the baseline map are same as those depicted in Fig. 4.B) Quantitative analysis of homotopic S1FP RSFC. 1-way repeated measures ANOVA did not reveal a significant effect of time, nor were there any differences between different time points. C) Ipsilateral RSFC between right S1FP and Right Motor cortex. 1-way repeated measures ANOVA did not reveal a significant effect of time, nor were there any differences between different time points. n.s.= not significant. Post hoc tests were performed as t-tests assuming unequal group variance and corrected for multiple comparisons using false discovery rate correction.

**Figure S4.**
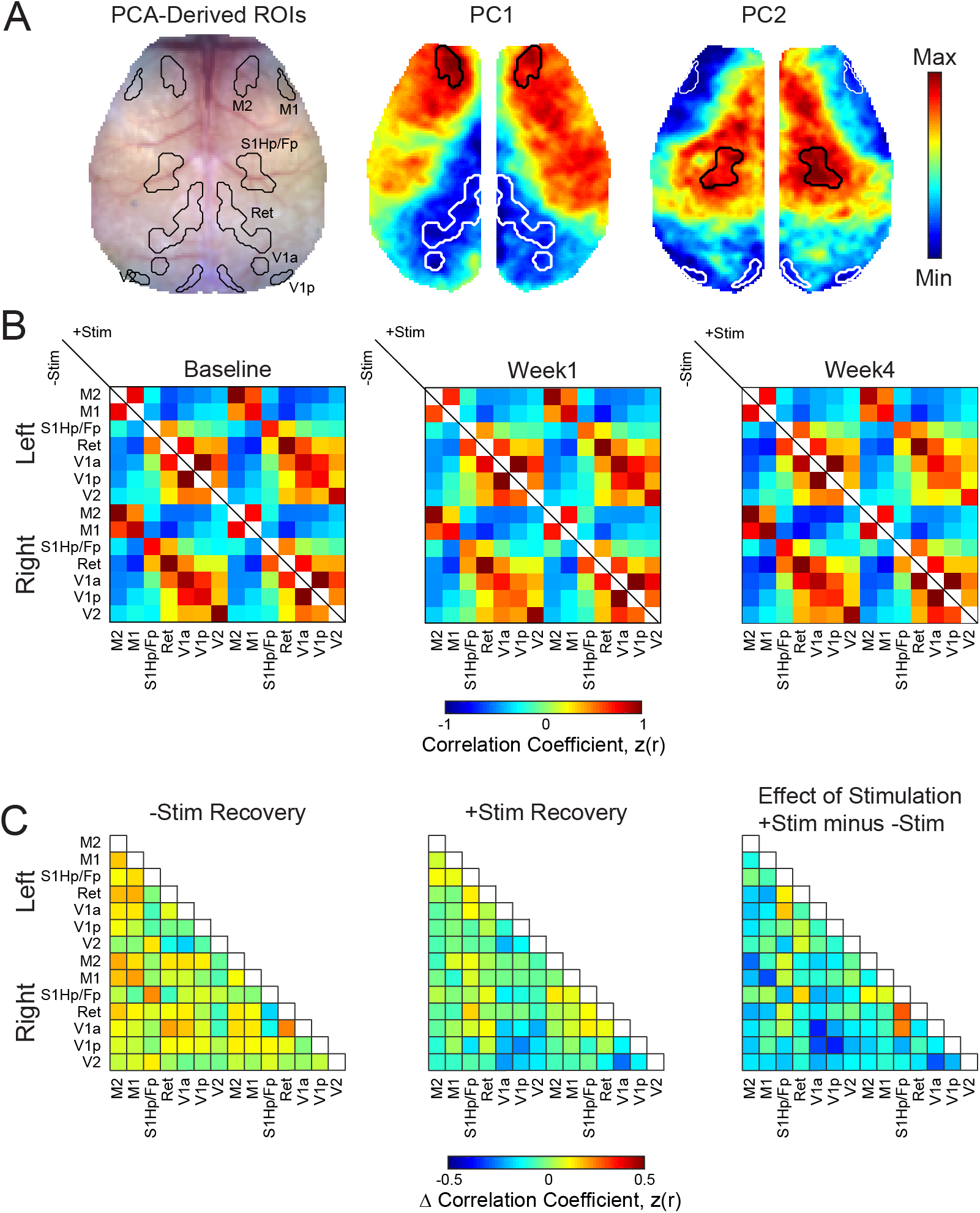
Regional RSFC between PCA-Derived ROIs and RSFC Recovery. A) White light image showing ROI contours and contours overlaid on respective PCs from which ROIs were determined. B) RSFC matrices for PCA-derived ROIs over time in –Stim (n=10) and +Stim (n=15) groups. Matrices are organized by functional assignment, then by hemisphere. RSFC of –Stim mice are reported in the lower triangle while that of +Stim mice in the upper triangle. RSFC is reported as Fisher z scores. C) Recovery of ROI-based RSFC. Recovery for each group was calculated as the magnitude of RSFC at Week 4 minus that at Week 1. The effect of stimulation was calculated as Recovery of the +Stim group minus recovery of –Stim group.

**Figure S5.**
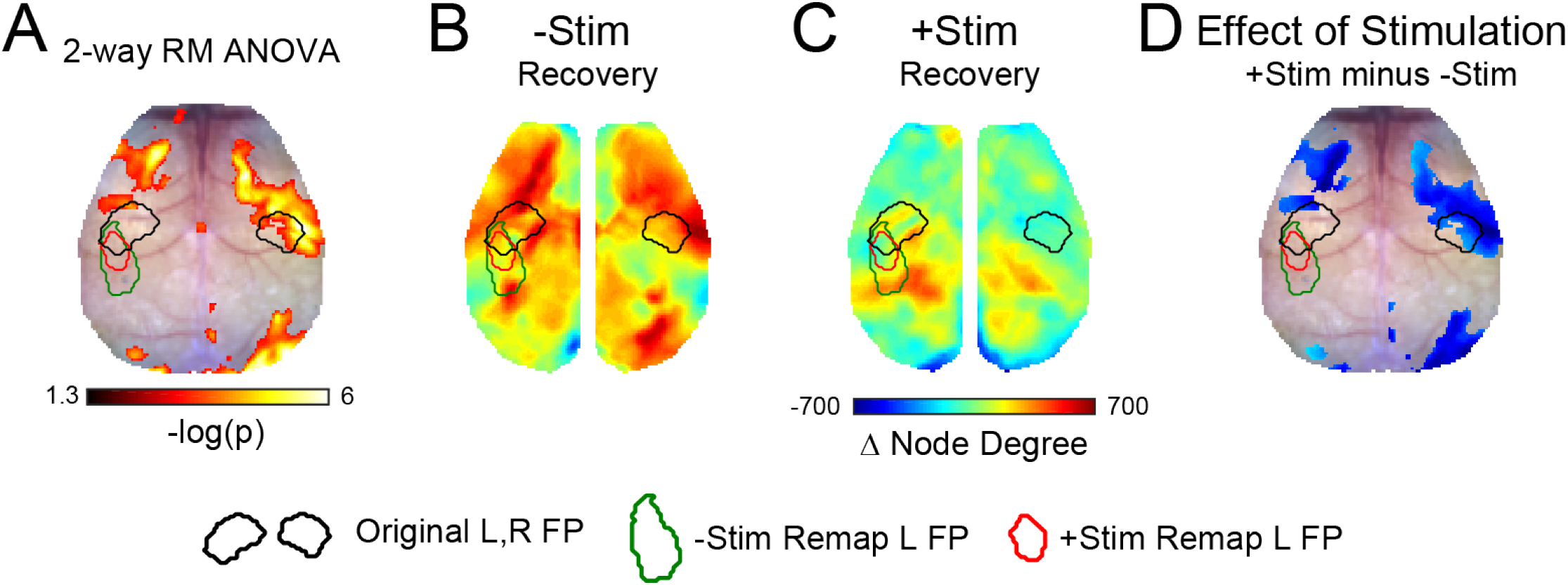
Effect of stimulation on node degree recovery. Repeated measures 2-way ANOVA computed at each pixel revealed a significant Group x Time interaction in parts of primary motor, large portions of secondary motor, right primary somatosensory forepaw and surrounding cortex, perilesional somatosensory cortex, and right posterior visual cortices. Cluster-corrected p-values shown on a –log scale. B) and C) Node degree recovery was determined for each group by subtracting maps reported in Figure 6 at Week 4 from those at Week 1 for both groups. D) Group-wise recovery differences were then calculated as +Stim recovery minus –Stim recovery. Blues indicate reduced node degree in +Stim mice compared to –Stim mice), and reds indicate increased node degree +Stim mice. Difference map is thresholded by the statistical map of Fig S5A. −Stim (n=10) and +Stim (n=15)

